# Comprehensive Variant Effect Map of Parkin-Mediated Mitophagy in Parkinson’s Disease

**DOI:** 10.64898/2026.02.09.704749

**Authors:** Erna Sól Sigmarsdóttir, Vasileios Voutsinos, Kristoffer Enøe Johansson, Isa Kristin Henrichs, Alissa Buhrmann, Kresten Lindorff-Larsen, Rasmus Hartmann-Petersen

**Affiliations:** Department of Biology, University of Copenhagen, Ole Maaløes Vej 5, 2200N Copenhagen, Denmark

**Keywords:** Deep mutational scanning, DMS, Parkinsons’s disease, PRKN, ubiquitin, mitochondria, PINK1/Parkin mitophagy, MAVE, ARPD, MAGIC, E3, variant classification

## Abstract

The development of Parkinson’s disease (PD) has a substantial genetic basis. Variants in the *PRKN* gene account for roughly half of autosomal-recessive PD cases, yet most variants remain classified as variants of uncertain significance. *PRKN* encodes the E3 ubiquitin-protein ligase, Parkin, that plays a key role in mitochondrial quality control by initiating mitophagy. In this work, we introduce a multiplexed mitophagy assay and measure the activity of more than 99% of all possible single–amino acid substitution and nonsense variants of Parkin. We also demonstrate that misfolded, rapidly degraded variants autonomously trigger Parkin-independent mitophagy. The obtained activity landscape closely reflects known structural and functional aspects of Parkin while revealing new insights into specific functional effects across the protein. This dataset near-perfectly distinguishes pathogenic from benign variants and surpasses both computational predictors and abundance-based metrics in pathogenicity assessment. Finally, the data pinpoint hypomorphic variants that could be amenable to rescue in future personalized therapeutic approaches for PD.

## Introduction

Mitochondria are vital organelles involved in critical cellular functions including ATP production, metabolism, and signaling^1^. During energy production, reactive oxygen species are generated in the mitochondria, leaving them highly susceptible to mutations in mitochondrial DNA, protein misfolding and dysfunction^2^. Accordingly, efficient regulation of mitochondrial dynamics and quality control is critical, including clearance of defective mitochondria through mitophagy^3^. During mitophagy, damaged mitochondria are engulfed by the autophagosome, which subsequently forms an acidic autolysosome, where hydrolases catalyze mitochondrial degradation^4^. The significance of mitophagy is highlighted by the wide range of conditions linked to its impairment, including heart failure, cancer and aging^1,3^. Mitochondrial dysfunction is also a major hallmark of Parkinson’s disease (PD)^5^ as well as other neurodegenerative disorders^1^.

The Parkin protein, encoded by the *PRKN* gene, is an E3 ubiquitin-protein ligase that plays a key role in mitochondrial quality control^6,7^. Parkin is made up of 465 amino acid residues and has five major domains: a ubiquitin-like (UBL) domain and four zinc-coordinating RING-like domains, namely RING0, RING1, IBR (in between RING) and RING2^8^. Additionally, Parkin contains an activation element (ACT) in the linker region between UBL and RING0, and a repressor element (REP) in the linker region between IBR and RING2^9,10^. Under basal physiological conditions, Parkin is kept in an auto-inhibited state where the UBL and REP occupy the E2 binding site in RING1, and RING0 blocks the catalytic cysteine residue at position 431 in RING2^11^. Upon depolarization of mitochondria, the serine/threonine kinase PINK1 phosphorylates ubiquitin at S65 and Parkin at S65 in the UBL domain^12^. Subsequently this leads to major structural rearrangements, including the exposure of the E2 binding site in RING1 and the binding of the UBL and ACT to RING0, which results in the release of the catalytic C431 residue in RING2^10^. In its activated state, Parkin ubiquitylates various proteins of the outer mitochondrial membrane (OMM), effectively tagging the mitochondrion for mitophagy^13^. Dysregulation of the PINK1/Parkin mitophagy pathway is closely linked to the pathogenesis of PD, although the details of how mitophagy protects against PD remain elusive^14^. Parkinson’s disease is characterized by the progressive loss of dopaminergic neurons in the *substantia nigra*, causing impairment of motor control, and affects about 2-3% of the global population over 65 years of age^5,15^. While most cases of PD are idiopathic (∼90%), a portion can be explained genetically with multiple known genetic risk factors^16–18^. Gene variants in *PRKN* have an especially strong link to early-onset PD. Further, homozygous or compound-heterozygous *PRKN* loss-of-function variants have been linked to the most common cause of autosomal-recessive PD (ARPD), accounting for about 50% of autosomal recessive parkinsonism in Europe^18,19^. In addition, pathogenic Parkin variants are commonly detected in patients with idiopathic PD^17^.

The pathogenic variants in *PRKN* are distributed across the gene and include missense, nonsense, and frameshift variants^20^. Nonsense and frameshift variants typically lead to nonsense-mediated mRNA decay or result in major changes to the encoded protein, and their consequences are usually deleterious and straightforward to predict. The effects of missense variants, on the other hand, can range from total loss of function, over neutral, to variants that are hyperactive^8,16,21–23^, thus making them more challenging to interpret and often leading them to be classified clinically as variants of uncertain significance (VUS)^8,24–26^. Accordingly, 71% of the 186 missense variants currently listed in ClinVar^20^ are categorized as VUS or have conflicting interpretations of pathogenicity. This large number of VUS limits the capacity to provide an accurate diagnosis and to offer effective genetic counselling to affected individuals and their families^8,24,27^.

To elucidate the functional effects of Parkin variants and uncover the molecular mechanisms by which missense mutations lead to pathogenicity, we previously applied the Variant Abundance by Massively Parallel Sequencing (VAMP-seq) technology^28^ to determine the cellular abundance of >99% of the possible single amino acid substitutions and nonsense Parkin variants^29^. This revealed that half of the pathogenic missense variants display a lowered cellular abundance, likely owing to a reduced structural stability and subsequent degradation^8,30^. Since the pathogenic variants can lead to loss of function without affecting structural stability^8,31^, abundance is on its own not a sufficiently powerful metric for classifying all variants^29^. Accordingly, for confident interpretation of variants observed in population or patient sequencing, and to gain a comprehensive understanding of the underlying molecular effects of Parkin variants, high throughput mapping of Parkin function is warranted.

Here, we present a functional assessment of 9,212 out of 9,300 (99.1%) possible single amino acid substitutions and nonsense Parkin variants using high throughput cell-based assays of Parkin-dependent mitophagy. We find that Parkin variant activity correlates with sequence conservation and captures most known pathogenic variants. The results highlight specific sites that are critical for Parkin activity and reveal that many pathogenic Parkin variants are hypomorphs and can therefore potentially be rescued by increasing Parkin stability or expression.

## Results

### Deep mutational scanning of Parkin variant activity

Building on our previous study on Parkin variant abundance^29^, we set out to perform a complementary high throughput assay focusing on Parkin variant activity. To this end we developed a multiplexed assay of variant effects (MAVE) using the mitochondria-targeted and pH-sensitive fluorescent protein, mt-mKeima, to quantify Parkin dependent mitophagy^32^. For this, we repurposed our previous cDNA library of Parkin variants^29^ to include the mt-mKeima reporter. This site-saturated and barcoded plasmid library of GFP-tagged Parkin variants, which allows co-expression of mt-mKeima, was integrated into a genomic landing pad in HEK293T cells via Bxb1 mediated recombination^33^. Before integration, the landing pad expresses a gene cassette encoding BFP, iCasp9, and a blasticidin resistance gene, separated by autocleaving 2A peptides. The iCasp9 expression allows induction of apoptosis in non-recombined cells with AP1903 (Rimiducid) treatment. With successful integration, the landing pad gene cassette is displaced and no longer expressed. In its place, a GFP-fused Parkin variant and mt-mKeima are produced from the same mRNA transcript (**Fig. 1A**). The vector does not include a promoter, ensuring that non-integrated plasmids are not expressed and a single Parkin variant is expressed per cell^33^. Under normal physiological conditions, the mitochondrial matrix is maintained at a near-neutral pH and mt-mKeima excitation occurs at 405 nm. During mitophagy, acidification shifts the excitation of mt-mKeima to 561 nm^32^. The ratiometric shift in mt-mKeima excitation can be used as a readout of mitophagy^21,23,29,32^, thus serving as a measure of Parkin activity (**Fig. 1A**).

**Figure 1.**
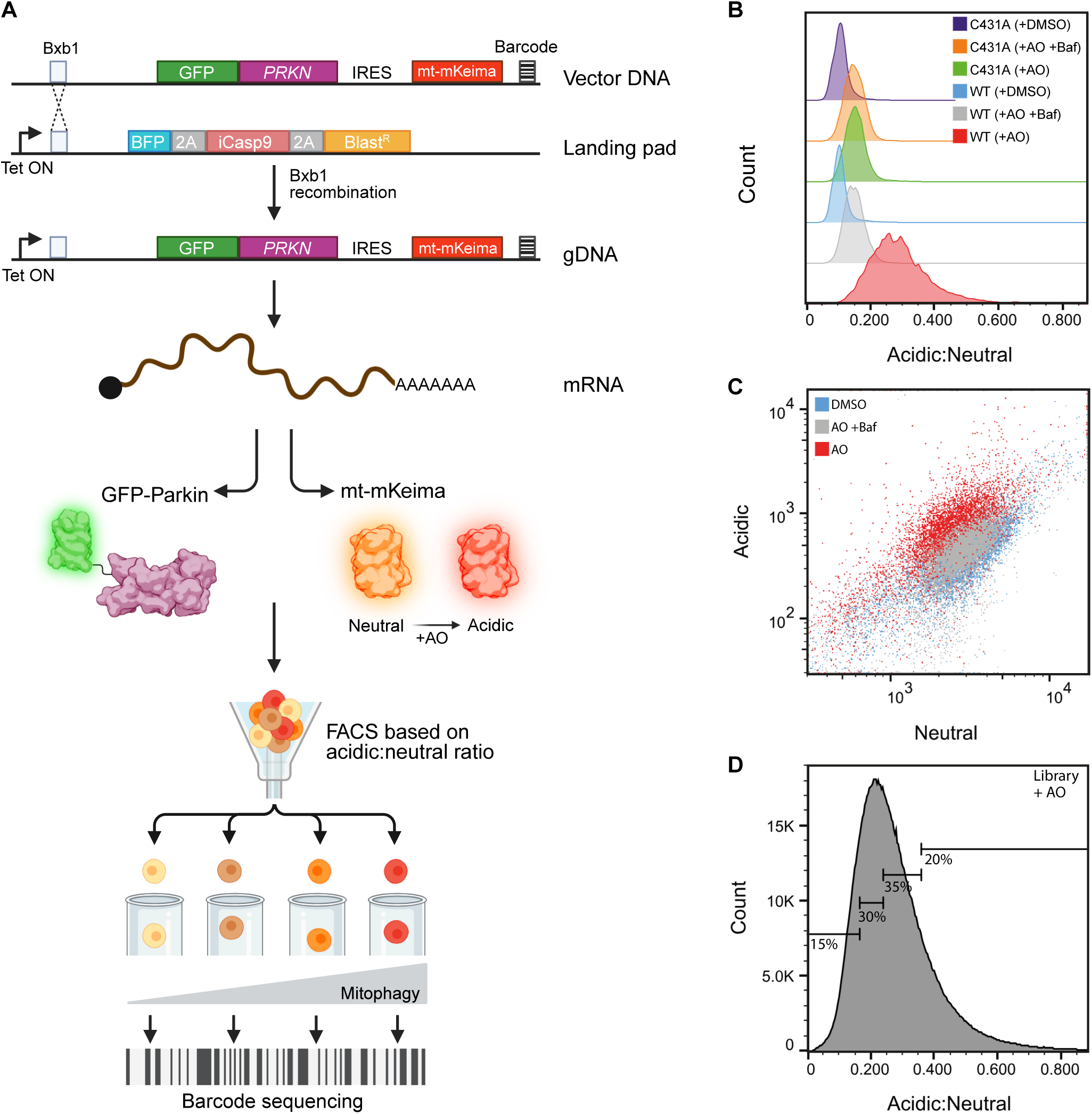
Parkin variant activity assay. (A) A schematic illustration of the expression system and Parkin variant activity assay. A *PRKN* variant library is integrated into the genome of HEK293T TetBxb1BPFiCasp9 cells through Bxb1 specific recombination at a genomic landing pad. Before recombination, the landing pad consists of BFP, iCasp9, and a blasticidin resistance gene (BlastR), separated by auto-cleaving 2A peptides^33^. Cells are transfected with a Bxb1 recombinase expression plasmid (not shown) and a vector encoding a GFP-fused Parkin variant and mt-mKeima separated by an internal ribosome entry site (IRES), a Bxb1 recombination site, and a barcode. With doxycycline treatment the Tet-On promoter is activated and a GFP-fused Parkin variant and mt-mKeima are produced from the same mRNA transcript. As Parkin activates mitophagy upon antimycin and oligomycin treatment (AO) the pH in the mitochondria drops, shifting the excitation of mt-mKeima^32^. The ratio of acidic:neutral mt-mKeima is thus used as a readout for mitophagy and Parkin activity. Fluorescence-activated cell sorting (FACS) is used to separate the Parkin variant library into four bins based on the acidic:neutral mt-mKeima ratio, and the barcodes are sequenced to identify the variants within each bin. Figure adapted from^29,36^. Created in BioRender. Hartmann-Petersen, R. (2026) https://BioRender.com/vdjorro. (B) Representative flow cytometry profiles showing the induction and suppression of mitophagy in cells expressing WT Parkin versus the catalytically dead variant Parkin C431A. AO induces mitophagy in cells expressing WT Parkin (red, n=9,702) compared to mock treated (DMSO) (blue, n=9,388). Treatment with bafilomycin suppresses mitophagy (AO+Baf) (grey, n=9,844). AO treatment barely affects cells expressing the catalytically dead C431A variant (green, n=9,487), compared to mock (vehicle control) treatment (purple, n=9,260). C431A with bafilomycin is included as a control (orange, n=9,625). (C) A representative flow cytometry scatterplot showing the AO triggered induction and bafilomycin (Baf) mediated suppression of mitophagy. HEK293T cells expressing the variant library were treated with AO for induction of mitophagy (red, n=1.46×10^6^), with AO and bafilomycin (AO+Baf) for suppression of mitophagy (grey, n=1.31×10^6^), or DMSO as a mock (vehicle control) treatment (blue, n=1.03×10^6^). The x-axis indicates the level of neutral mt-mKeima and the y-axis the level of acidic mt-mKeima. (D) FACS sorting strategy. Cells expressing the Parkin variant library were sorted into four bins based on their acidic:neutral mt-mKeima ratio. The first bin was set to contain 15% of the total cell population with the lowest ratio. The second bin was set to contain 30% of the total population. The third bin was set to contain 35% of the total population and the fourth bin was set to contain 20% of the total population with the highest ratio. The figure shows the profile and gating for the AO treated population. Data for the DMSO treated cells are included in the supplementary material (**Supplementary** Fig. 2).

To test this system, we first used flow cytometry to compare the acidic:neutral mt-mKeima signal of cells expressing either wild-type (WT) Parkin or the catalytically dead C431A variant^11^. Treatment with antimycin A and oligomycin A (AO), which together block oxidative phosphorylation and result in mitochondrial depolarization, robustly induced mitophagy in cells expressing WT Parkin, but only mildly in cells expressing the C431A variant (**Fig. 1B**). The shift in signal was reversed by blocking lysosomal acidification with bafilomycin A1, confirming that the observed increase in acidic mt-mKeima signal upon AO treatment is due to mitophagy. Since the catalytically dead variant failed to induce mitophagy, we conclude that any contributions from endogenous Parkin, or Parkin independent mitophagy pathways, are minimal for these variants. HEK293T cells express endogenous Parkin, but at very low levels, and overexpression is generally required for functional mitophagy assays^34,35^. In agreement with this, we are unable to detect endogenous Parkin in the landing pad cell line by western blotting^29^, and by RNA sequencing the endogenous *PRKN* level falls within the noise of the experiment^36^. To further validate the system, we used fluorescence microscopy to examine the localization of WT Parkin and the C431A variant. Upon AO treatment, WT Parkin translocated to the mitochondria, as indicated by co-localization of the GFP-tagged protein with MitoTracker staining of the mitochondria. In contrast, the catalytically dead C431A Parkin remained diffusely localized in the cytosol (**Supplementary Fig. 1**).

Based on the above data, we deemed the separation in mt-mKeima signal between WT and C431A cells provided a sufficient dynamic range for high throughput screening and proceeded with integrating the site saturated Parkin library into the landing pad. Flow cytometry of library transfected cells revealed the expected shift in acidic:neutral mt-mKeima signal upon treatment with AO, which was reversed by bafilomycin (**Fig. 1C**). We then proceeded to sort the cells into four bins by fluorescence-activated cell sorting (FACS). As we wanted to ensure a high resolution for both the inactive and the hyperactive variants, we designed the bins so that the first, low activity bin covered 15% of the total cell population with the lowest fluorescence ratio, the second bin covered 30%, the third bin covered 35%, and the fourth, high activity bin, covered 20% of the total cell population (**Fig. 1D, Supplementary Fig. 2**). In parallel to the AO treated cells, we also sorted DMSO treated cells using the same bin setup, to assess background signal. We used DMSO rather than untreated cells, as AO was dissolved in DMSO, and DMSO alone induces low levels of mitophagy (**Supplementary Fig. 3**).

### A functional map of Parkin variants

After sorting the cells into the four bins based on the acidic:neutral mt-mKeima signal (**Fig. 1D**), we isolated genomic DNA and sequenced the barcodes to determine the frequency of each variant within each of the four bins. This allowed us to calculate the average bin that each variant occupies under both DMSO (**Supplementary Fig. 4**) and AO treatment (**Supplementary Fig. 5**). From these data we calculated an activity score that represents the AO-induced mitophagy (see Methods) for each of the variants in the library (**Supplementary Fig. 6**). This score ranges from about 0 to 1, with 1 representing a mitophagy similar to the WT Parkin, while a score of 0 is indicative of low mitophagy similar to the catalytically dead C431A variant. These scores and their standard deviations were derived from three biological replicates (separate library transfections) and show strong consistency across replicate experiments, with Pearson’s correlation coefficients ranging from 0.82 to 0.85 for the screens with AO treatment and from 0.52 to 0.62 for the DMSO treated screens. The relatively poor correlation for the DMSO treated screens was expected given the low signal without AO-induced mitophagy, which results in a lower signal to noise ratio (**Supplementary Fig. 7**). The dataset includes activity scores for 9,213 out of 9,300 possible variants (calculated as: 465 residues × 19 amino acid substitutions per position, 465-1 positions for early stop codons, and 1 WT) representing a coverage of 99.1% (**Fig. 2A**). All data are included in the supplementary data file (**Supplementary Data File**). We also obtained activity scores for synonymous (silent) variants in 419 out of 465 positions (90.1%) (**Supplementary Data File**). At positions 1, 344, and 345 the majority of substitutions were missing, primarily due to failures in library synthesis and cloning. The activity score distribution featured one large peak centered around a score of 1, overlapping with the synonymous variants and the score of WT Parkin, and smaller “shoulder” at a score of about 0, overlapping with the catalytically dead C431A and nonsense variants (**Fig. 2B**).

**Figure 2.**
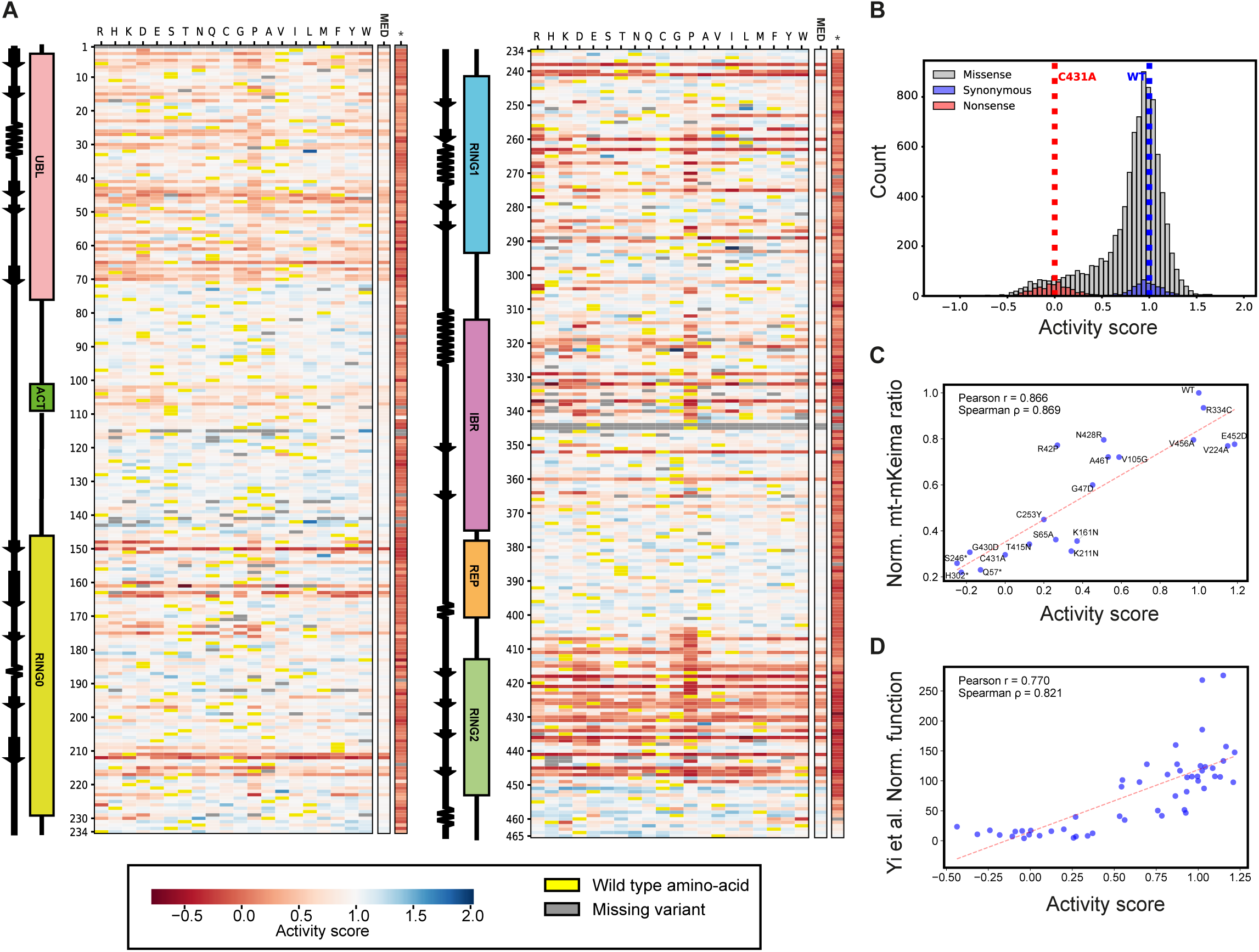
Variant effect map of Parkin activity. (A) Heat-map of Parkin activity scores. The scores included in the map are the ones with a standard deviation (SD) lower than the “knee” of the SD distribution. Median scores (MED) were only calculated from the included data. The asterisks (*) indicate nonsense variants. The horizontal axis represents each possible amino acid. The vertical axis shows the positions in Parkin. The color scale is shown at the bottom of the map: WT-like activity (white); lower than WT activity (red); higher than WT activity (blue). The WT (reference) amino acids are marked yellow and missing variants in grey. The linear organization of Parkin domains and the secondary structure elements across the sequence are included. (B) Histogram distribution of the Parkin activity scores for all missense (grey), synonymous (blue) and nonsense (red) variants. The scores for the controls, C431A (red) and WT (blue), are indicated with dotted lines. (C) Scatterplot comparing the Parkin activity scores from the screen (x-axis) with the low throughput acidic:neutral mt-mKeima ratio measurements of selected AO treated variants after subtraction of the equivalent measurements under DMSO treatment and normalization to the WT (y-axis) (for WT and C431A n = 8, for T415N and G430D n = 6, for the nonsense variants n = 2, for the rest of the variants n = 3). (D) Scatterplot comparing the Parkin activity scores (x-axis) with the normalized function as obtained by Yi *et al*. (2019)^21^ (y-axis).

We validated the results of the multiplexed assay using low throughput flow cytometry measurements of cells individually transfected with 20 Parkin variants (**Fig. 2C, Supplementary Table 1**), and with data from Yi *et. al.* (2019)^21^ on activity of 52 variants determined in U2OS cells, with good correlations (**Fig. 2D**) (Pearson r = 0.87 and Pearson r = 0.77 respectively). However, in the comparison with the Yi *et. al.* (2019) data we noticed a loss of resolution for hyperactivity, which is likely due to our assay being based on the acidic:neutral mt-mKeima ratio instead of the percentage of cells with acidic mt-mKeima above a certain threshold, which only considers the cells with the most activity. In addition, because our setup relies on overexpression, this may contribute to the reduced sensitivity for hyperactive variants, and account for the absence of detectable hyperactivity of the well-characterized W403A variant^9,21–23,37–39^ and other residues at the RING0:RING2 interface (**Supplementary Table 2-4**). Nonetheless, we do identify multiple variants that have previously been reported to exhibit hyperactivity, including Y143D^23^, Y143E^23^, A401K^23^, F463Y^23^, V224A^21,22^, R256C^21,22^, and M458L^21,22^ (**Fig. 2A**). To avoid overinterpreting uncertain measurements, we applied a filter to remove variants with the largest uncertainties (**Supplementary Fig. 8A**), so that the final set of variants in our Parkin activity heat-map includes 9,048 of the 9,213 (98.2%) measured variant scores. Indeed, this filter removed a large portion of variants with extreme activity scores (**Supplementary Fig. 8B**).

Our mutational map showed, as expected, that many of the most sensitive positions were localized to the catalytic RING2 domain (**Fig. 2A**), but several other sites also appeared critical, including the PINK1 phosphorylation site at S65, where only a substitution to threonine is tolerated and the phosphomimetic aspartate and glutamate appeared to display partial activity (**Fig. 2A**, **Table 1**). An important step in Parkin activation is the binding of the phosphorylated UBL (pUBL) domain to RING0. This binding leads to release of RING2 and frees the active site^10,40^. In our map, residues at positions I44–G47 around the hydrophobic patch in the UBL domain appeared critical (**Fig. 2A, Supplementary Table 2**). This is presumably due to their position at the binding interface of pUBL and RING0. Within this interface, a positively charged pocket in RING0 binds pUBL during Parkin activation. The key residues of this phospho-acceptor pocket are K161, R163, and K211^10^, and their importance is also evident in the mutational map (**Fig. 2A**, **Table 1**). Other residues (e.g. V70, V164, and T173–L176) at this interface also appear important, although to a lesser extent (**Fig. 2A, Supplementary Tables 2 and 3**).

**Table 1.**
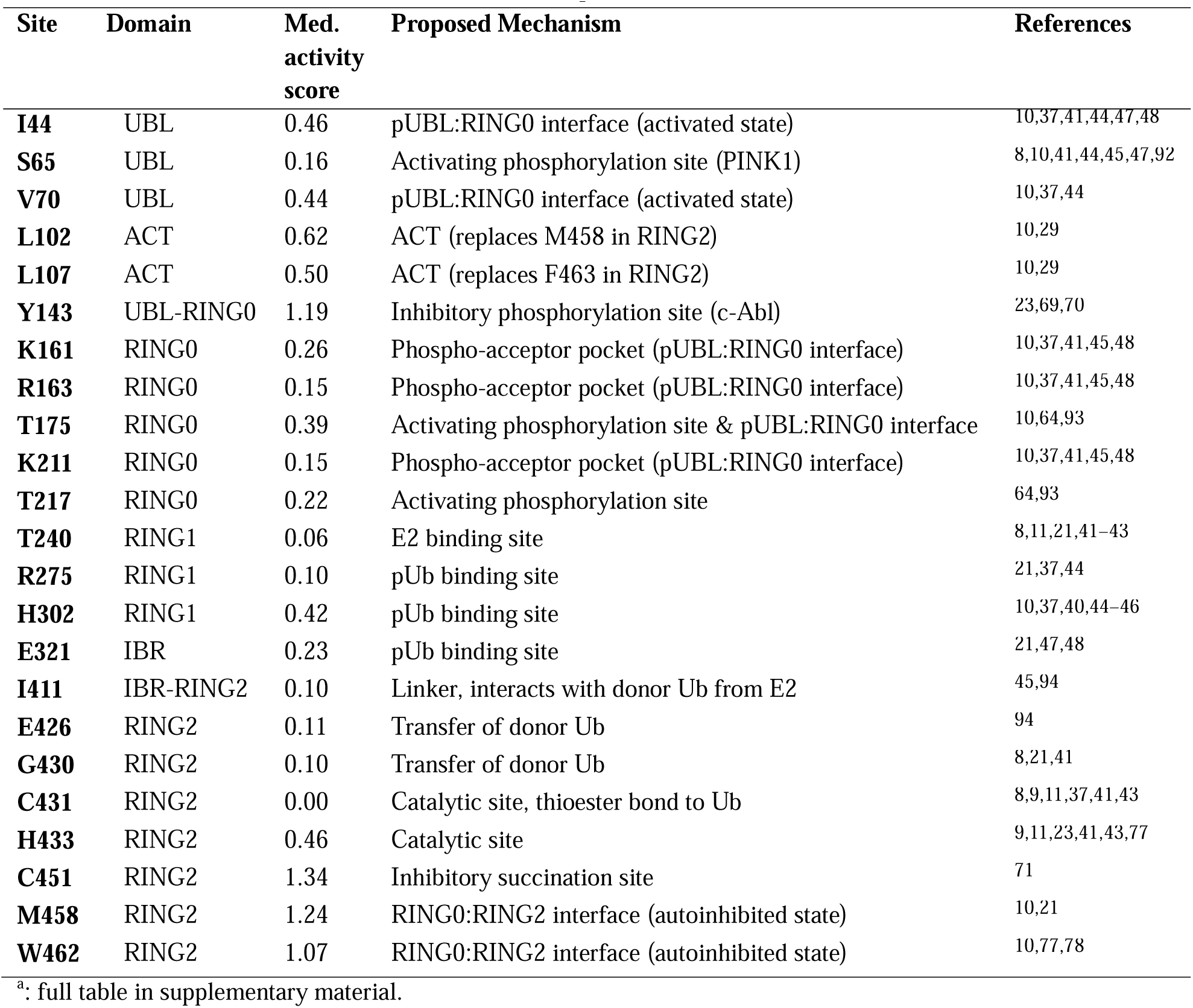
Selected notable positions in Parkina.

Another step in Parkin activation is the binding of ACT to RING0, which further helps the dissociation of RING2 from RING0. In particular, the three residues, L102, V105, and L107, in ACT directly displace the RING2 residues M458, W462, and F463 in the RING0:RING2 interface^10^. In agreement with this, the mutational map shows that these ACT residues are important, although some substitutions, mostly to other hydrophobic amino acids, are tolerated (**Fig. 2A**, **Table 1**).

As expected, substitutions around residues T240 ^8,11,21,41–43^ and Y267^41^ were not tolerated, which is likely attributed to their role in E2 binding (**Fig. 2A**, **Table 1**). Likewise, substitutions around R275^21,37,44^, H302^10,37,40,44–46^, and E321^21,47,48^, reported to be involved in phospho-ubiquitin (pUb) binding, were not tolerated (**Fig. 2A**, **Table 1**). At position K151^10,40,44,45^ substitutions to hydrophobic and negatively charged residues affected mitophagy levels, while others were tolerated. Surprisingly, substitutions of R305^10,37,44,45^, also frequently reported as important for pUb binding, only resulted in minimal reduction of mitophagy (**Fig. 2A, Supplementary Table 4**).

The mutational map also shows the importance of the zinc-coordinating residues in Parkin. Four domains in Parkin, RING0, RING1, IBR, and RING2, each bind to two Zn^2+^ ions through conserved cysteine and histidine residues^9^. Accordingly, many of the zinc-coordinating residues, especially those in RING1, IBR, and RING2, appear important for Parkin activity (**Fig. 2A, Supplementary Table 2**).

Beyond these examples we highlight additional positions (**Table 1**, **Supplementary Table 2**), including some that cannot be readily explained by the current understanding of Parkin structure and function (**Supplementary Table 3**).

Overall, the determined variant effects on Parkin function aligns well with established structural and functional hotspots of the protein, while at the same time offering a comprehensive overview across all positions.

### Low abundance protein variants induce Parkin-independent mitophagy

We observed an overall increased mitophagy for variants that we previously^29^ found to be of low abundance, even in the absence of AO treatment (**Supplementary Fig. 9**). These variants mostly include misfolded missense and nonsense variants, but also nonsense variants within the disordered linker downstream of the UBL domain, which are predicted to produce C-degrons using the protein abundance predictor (PAP)^49^ (**Supplementary Fig. 10**). Nonsense Parkin variants tested in low throughput also displayed increased levels of background mitophagy in DMSO (without AO) compared to wildtype (**Supplementary Fig. 11A**). These observations suggested that expression of some misfolding-, degradation- or aggregation-prone proteins might induce mitophagy independent of the Parkin activity. To investigate this further we used the landing pad system to express the non-mitophagy related protein ASPA and the low abundance and misfolded C152W ASPA variant^36,50^. Comparing the mitophagy activity based on the mt-mKeima reporter revealed that cells expressing the unstable C152W ASPA variant exhibited increased levels of mitophagy compared to the WT ASPA, without addition of AO (**Supplementary Fig. 11B**). The same was observed for the rapidly degraded R42P Parkin variant, which, when overexpressed, has higher levels of background mitophagy compared to cells expressing WT Parkin. A R42P,C431A double mutant, which additionally is inactive, showed the same increased background mitophagy as the R42P single variant (**Supplementary Fig. 11C**). These results show that misfolded or rapidly degraded overexpressed proteins induce mitophagy, independently of Parkin activity, presumably via the mitochondria as guardian in cytosol (MAGIC) pathway^51^ or perhaps simply by draining the cells of energy, due to increased levels of protein degradation.

### Parkin activity in relation to its structure and to computational predictors

To further analyze the functional effects of Parkin variants, we noticed that at many positions the tolerance to substitutions appear to depend strongly on the position in the protein (**Fig. 2A**) with position median scores explaining 61% of the variance of all missense variants (Pearson 0.78). The AlphaFold2 predicted Parkin structure (AF-O60260-F1), which resembles the crystal structure of Parkin in the closed conformation^9,10,44^, was colored with the median mitophagy score for each position (**Fig. 3A**). This revealed that most sensitive positions were buried and/or clustered in the UBL domain or in RING2 near the active site. There are only few positions where many variants are hyperactive, and they are primarily positioned close to or within the RING2 domain.

**Figure 3.**
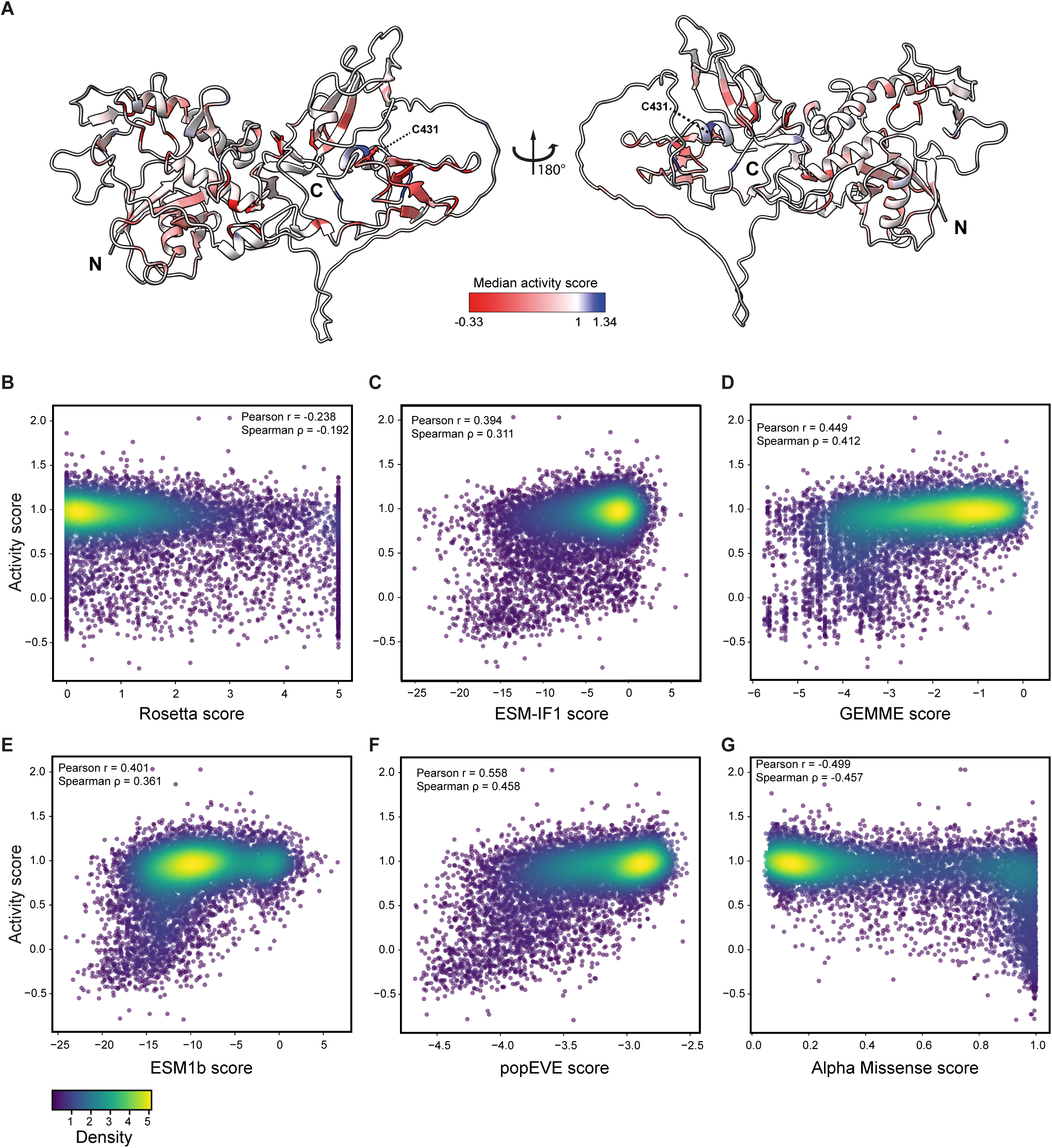
Comparisons of the variant effects with the computational predictions. (A) Cartoon representation of the Parkin structure (AF-O60260-F1) as predicted by AlphaFold2. The structure is colored by the median activity scores. The color scale is shown in the middle. The N-and C- termini are indicated and the catalytic C431 is highlighted in stick representation. (B-G) Scatterplots correlating Parkin activity scores with variant effect predictions based on (B) Rosetta, (C) ESM-IF1, (D) GEMME, (E) ESM1b, (F) popEVE, and (G) AlphaMissense.

Next, we compared our results with computational variant effect predictions (**Fig. 3B-G**). In general, the experimental results were overall poorly captured by protein stability predictions such as with Rosetta^52^, which estimate the change in thermodynamic folding stability compared to WT Parkin, and the inverse protein-folding language model ESM-IF1^53^ (Spearman’s ρ = - 0.192 and 0.361, respectively) (**Fig. 3B,C**). This was expected given that many variants may affect the activity without overly destabilizing the protein structure. Accordingly, GEMME^54^, which predicts the evolutionary “cost” of introducing substitutions based on multiple sequence alignments of Parkin orthologues, revealed moderate correlation with the activity scores of our screen (Spearman’s ρ = 0.412) (**Fig. 3D**).

Similarly, we used variant effect predictors ESM1b^55^, popEVE^56,57^ that are in part based on protein language models, and AlphaMissense^58^ which is based on the AlphaFold2 model for protein structure and sequence. These exhibited the best correlations but clearly did not capture all the observed effects (**Fig. 3E-G**). We speculate that part of the disagreement is due to the overexpression of *PRKN* variants in our system. Thus, variants that are hypomorphs may, when overexpressed, appear functional in our assay, even though they are predicted as unstable and potentially pathogenic (**Supplementary Fig. 12)**. We find many such low abundance variants with normal activity in RING1, RING0 and UBL domains (**Supplementary Fig. 13**), all of which are involved in the repression of Parkin activity, likely indicating that even though they are low in abundance they are constitutively active, therefore scoring high in our activity screen.

### Combination of Parkin activity and abundance can predict pathogenicity

Previously we found that Parkin variant abundance on its own is not sufficient as a metric to discriminate between all pathogenic and benign missense variants^29^. To probe this further, we compared the abundance scores with the activity scores (**Fig. 4A**). This revealed that many variants, including most variants classified as benign or likely benign in ClinVar^20^ (downloaded July 2023) and MDSGene^59^, displayed WT-like abundance and activity scores (close to 1), while most pathogenic or likely pathogenic variants showed reduced abundance and/or activity scores. We then applied Youden’s J statistic^60^ to define the thresholds for the activity (0.371) and abundance (0.639) scores that separate benign/likely benign variants from pathogenic/likely pathogenic variants. In turn, this allowed us to separate the variants into four groups: WT-like stable and active variants (top right quadrant), low abundance but active hypomorphs (top left), low abundance and inactive variants (bottom left), and stable but inactive catalytic variants (bottom right) (**Fig. 4A**). This revealed that in contrast to the abundance scores, the activity scores almost perfectly separate the benign/likely benign from the pathogenic/likely pathogenic variants (**Fig. 4A,B**). Four variants: the pathogenic R33Q, the likely pathogenic G328E, and the low abundance V56E and G284V, score above the activity threshold despite their pathogenic status. On the other hand, all benign/likely benign variants were properly assigned to the active quadrants. When taking into account both the abundance and activity scores, only R33Q and G328E are incorrectly categorized.

**Figure 4.**
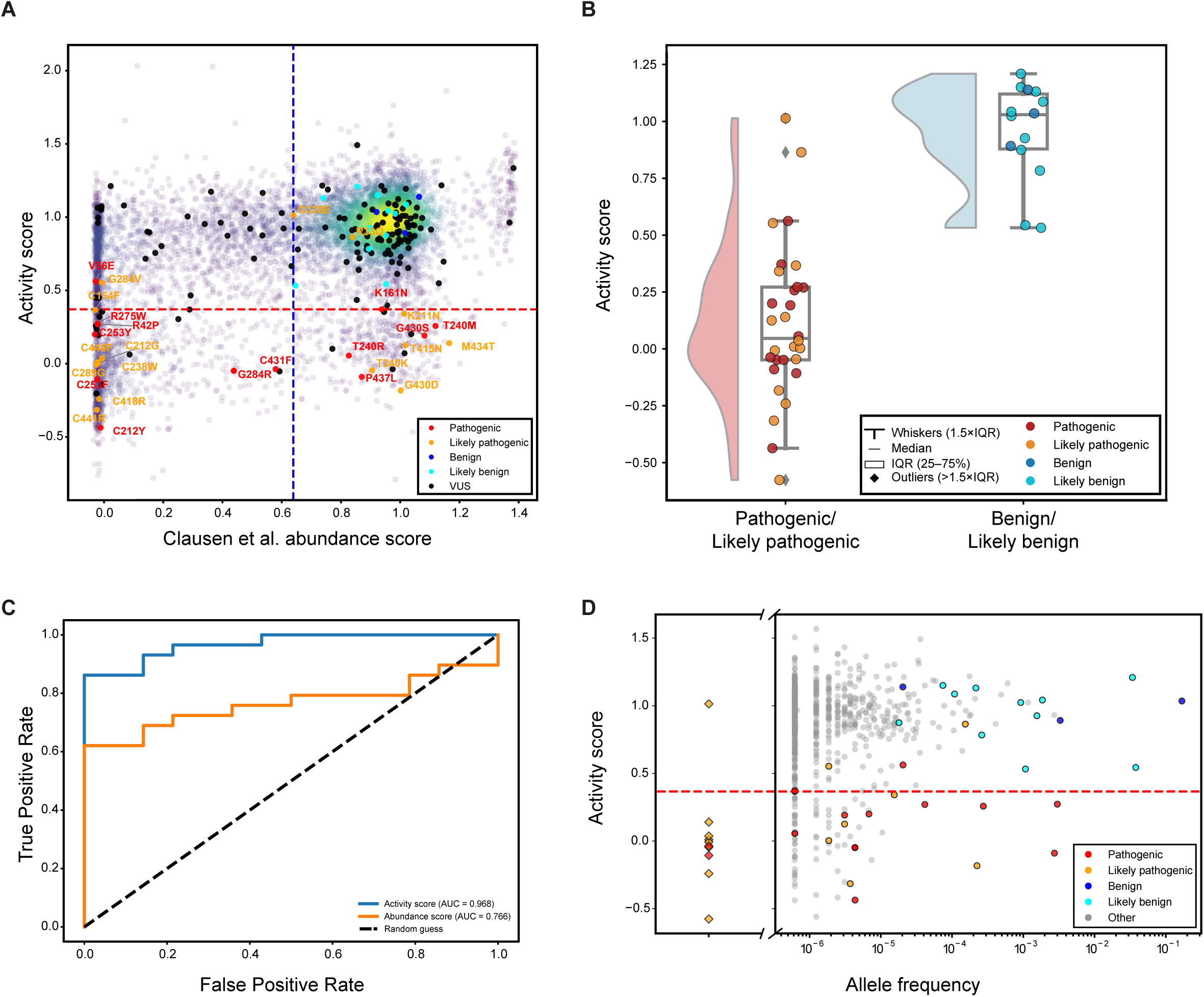
Applying the Parkin MAVE scores for analyzing pathogenic variants. (A) Scatterplot of the Parkin abundance scores as obtained from VAMP-seq^29^ (x-axis) with the Parkin activity scores (y-axis). Known pathogenic (red), likely pathogenic (orange), benign (blue), likely benign (cyan) and variants of unknown significance (VUS, black) are highlighted. The pathogenic/likely pathogenic variants are annotated. The dotted lines mark the determined thresholds based on Youden’s J statistic for activity (red) and abundance scores (blue). (B) Raincloud plots illustrating the activity score distributions of pathogenic/likely pathogenic (red/orange) and benign/likely benign (blue/cyan) Parkin variants as reported in ClinVar. Dots are scattered on the horizontal axis to limit overlap. Boxplots are defined as indicated in the insert. IQR: interquartile range. (C) ROC curves for activity scores (blue) and abundance scores (orange). The black dotted diagonal line marks a random guess. (D) Scatterplot comparing the Parkin activity scores with the Parkin allele frequency of pathogenic (red), likely pathogenic (orange), benign (blue), likely benign (cyan) and other Parkin variants (grey) annotated in the Genome Aggregation Database (gnomAD). Diamonds represent variants without known allele frequency. The red dotted line marks the threshold based on Youden’s J statistic for activity scores.

Further analysis using the Receiver Operating Characteristic (ROC) revealed that the activity scores reach an area under the curve (AUC) of 0.968, while the abundance scores achieve an AUC of 0.766 (**Fig. 4C**). The activity scores were also superior to the computational predictors (**Supplementary Fig. 14**). In the case of AlphaMissense (AM) this was due to three variants (T240M, G430S and P437L) (**Supplementary Fig. 15**), all of which display high abundance and two positioned near the active site C431. Interestingly, the inactive and low AM scoring variants mostly consist of variants that are close to the active site or important to the derepression of Parkin, with only few low abundance ones (**Supplementary Data File**).

Finally, we compared the activity scores with the allele frequency of variants reported in gnomAD^61^. As expected, this revealed that most of the variants that are most common in the sequenced population are benign or likely benign with activities above the threshold (**Fig. 4D)**, while many rare variants display strongly reduced activity. The data presented here indicate that these are likely to be pathogenic.

## Discussion

Deciphering how genetic variation alters protein function and contributes to pathogenicity, remains an enormous and pressing challenge^27^. Based on the size of the human genome, the mutation rate, and the global population, every non-lethal single nucleotide variant should exist in multiple living individuals^62^. This warrants assessment of not only observed VUS but essentially all variants, including those not yet documented by population sequencing^63^. Our previous deep mutational scan of Parkin protein variant abundance^29^ provided large-scale information on Parkin variant degradation but also showed that abundance alone was not sufficient to determine pathogenicity. Thus, because genetic variation can influence protein function and stability in multiple ways, a single parameter is often insufficient to capture the complete picture of variant effects.

Here, we multiplexed a mitophagy assay to determine the activity of 99.1% of all possible Parkin missense and nonsense variants. The observed variant effects closely match the current understanding of Parkin structure and function and provide a comprehensive functional dataset to support future mechanistic studies and genotype-phenotype correlations. We also identified several new positions of functional importance that cannot yet be fully explained and warrant further investigations. For instance, phosphorylation of T175 and T217 by PINK1 has been reported to induce translocation of Parkin to mitochondria^64^. This finding has been controversial, with other researchers not being able to detect PINK1-mediated phosphorylation at these sites, and data showing that these residues are not required for mitophagy^65–67^. Our map shows that several substitutions at these positions lead to a reduced mitophagy, suggesting that these threonine residues are functionally important, although not necessarily due to phosphorylation.

Surprisingly, substitutions of S108/109/110, which are ULK1 phosphorylation sites involved in early response to mitochondrial stress and activation of Parkin^68^, did not seem to drastically affect mitophagy. This might be because substitution of these residues to alanine has been shown to primarily delay, rather than prevent, Parkin recruitment to mitochondria^68^. Our mitophagy measurements were performed after 4 hours of AO treatment, which may have masked any early kinetic defects caused by these mutations. Nevertheless, for all three positions, substitution of serine with threonine was among the best-tolerated changes, underscoring the functional importance of these phosphorylation sites.

In addition to these phosphorylation sites and the well-characterized PINK1 phosphorylation site at S65, Parkin has also been reported to be phosphorylated at Y143 by the tyrosine kinase c-Abl^69,70^. This modification is inhibitory to Parkin activity, consistent with our observation that most substitutions at and near this site result in hyperactivity.

At position C451, we observed that almost every substitution leads to hyperactivity, with a median activity score of 1.34 for this site. Consistent with the hyperactivity, succination of C451 has been shown to inhibit Parkin activity^71^. C323 has also been reported as a site of succination^71^, but we did not observe any pronounced effects for substitutions at this position, which could be linked to the reported S-nitrosylation at this site, which can be either activating or inactivating^72–75^.

Some studies have proposed that Parkin contains a catalytic triad composed of residues C431, H433, and E444^11,41,76^. However, others have reported a C431 and H433 catalytic dyad^43,77^. Our data support this latter view, showing that most substitutions at E444 are tolerated.

Intriguingly, several naturally occurring hyperactive Parkin variants have been described^21,22^ ^9,23,46^. We identify several variants that lead to increased mitophagy, primarily at positions that make up the RING0:RING2 binding interface in the autoinhibited conformation^10,21,77,78^. Substitutions at these positions likely disrupt the binding of RING0 to RING2 and lead to release of the catalytic site, leaving the enzyme constitutively active. Similarly, substitutions of three of the Zn-coordinating cystine residues in RING0, C166, C169 and C196, surprisingly did not lead to loss of mitophagy. We hypothesize that when Zn-coordination is lost, the whole RING0 domain unfolds to a state where it cannot bind RING2 to stabilize the autoinhibited conformation and therefore leads to Parkin activity. Thus, inhibiting RING0:RING2 binding could have therapeutic potential to increase Parkin function in PD patients.

From our data we observed a correlation between low abundance variants and increased mitophagy without AO induction. This mitophagy appeared to be Parkin-independent and related to increased degradation. In 2017, Ruan *et al*. described a degradation pathway for cytosolic protein aggregates in budding yeast and human cells, termed MAGIC (mitochondria as guardian in cytosol)^51^, where proteins are imported into the mitochondrial matrix and degraded by mitochondrial proteases^51^. In addition, several studies in human cells have reported that misfolded proteins are imported into mitochondria, triggering mitochondrial stress and ultimately leading to a reduced mitochondrial content^47,51,79–83^. Thus, mitophagy possibly serves as a fail-safe mechanism to limit buildup of misfolded proteins, a notion supported by the observation that in human dopaminergic neurons α-synuclein aggregates physically interact with mitochondria leading to their degradation via Parkin-independent mitophagy^83^. We found that structurally unstable, low abundance Parkin missense and nonsense variants, as well as the unrelated misfolded ASPA C152W protein, all autonomously induce mitophagy without treatment with AO. Based on these observations, we hypothesize that these overexpressed and unstable proteins are targeted by the MAGIC pathway, thereby triggering mitophagy. In addition to this, unstable variants of the PD associated protein DJ-1 have also been identified as potential MAGIC substrates^84^. However, the underlying mechanism and its relevance for endogenous proteins *in vivo* remains to be determined. However, since predicted C-degron forming nonsense variants in the UBL-RING0 linker region, which we would expect to be structurally stable, also induce mitophagy, the observed effect could simply be a consequence of saturation of the proteasome. This would lead to higher levels of misfolded proteins that sequester cellular chaperones and/or depletion of ATP. Consequently, mitochondria may be indirectly damaged, given that both chaperones and ATP are important in the translocation of mitochondrial proteins into mitochondria^85,86^.

Beyond these mechanistic insights, our data contribute to the interpretation and classification of the Parkin missense variants currently recorded in ClinVar^20^ and MDSGene^59^, but also for variants that have not yet been observed in the population. We demonstrated by ROC analysis that the activity screen alone outperforms even the best performing predictor AM. We found that a lot of the high abundance variants in the vicinity of the active site or participating in the activation of Parkin are given a low AM score (indicating WT-like behavior). This suggests that AM might misclassify inactivating missense mutations, without a common underlying mechanism like lowering of protein abundance, thus underlining the importance of experimental verification of such predictors. By combining our overexpression-based mitophagy assay with the previous abundance data, based on the same *PRKN* cDNAs expressed in the same cell line, we also probed the molecular mechanisms leading to loss of function and e.g. identify hypomorphic variants such as R42P that remain functional but are pathogenic due to insufficient abundance. Such variants could potentially be rescued by therapeutics that increase variant abundance, e.g. through structural stabilization by small molecular folding correctors or deubiquitylating targeted chimeras (DUBTACs), such as those developed to p53 variants^87^.

While preparing this manuscript, we learned that the Kudla group was working on a related project^88^. In that work, pUb accumulation was measured for about 9200 Parkin variants. Overall, our mt-mKeima-based activity scores correlate well (Pearson r=0.77) with the pUb scores, demonstrating generally high concordance between two MAVEs performed using independent library generation and different functional readouts. The Kudla lab also generated an abundance map that correlates reasonably well (Pearson r=0.71) with our previously published VAMP-seq data^29^. When we examined how well the four maps can be used to separate pathogenic and benign variants we thus find overall agreement (AUC 0.97 and 0.94 for our mt-Keima and the Kudla group’s pUb-based functional maps, respectively, and AUC 0.77 and 0.64 for our VAMP-seq and the Kudla group’s antibody-based abundance map, respectively). The few differences between the two functional datasets may be linked to the different readouts, different duration and concentration of AO treatment, as well as different cell lines (HEK293T and HeLa), which could have different levels of background mitophagy. Nevertheless, the independently generated maps lead to comparable conclusions, demonstrating how MAVEs have become a robust technique to quantify the functional consequences of missense variants on protein function and abundance.

In conclusion, here we provide an example of how integrating functional and abundance data allows more accurate separation of benign and pathogenic variants. Beyond variant classification, these results also provide mechanistic insight into how specific Parkin variants alter protein behavior, supporting future personalized medicine and genetic counseling in relation to Parkinson’s disease.

## Methods

### Plasmids and library preparations

Integration plasmids of the single Parkin variants, for study in low throughput, were purchased from Genscript and were based on the VAMP-seq vector^29^, except for the mCherry, which was replaced by mKeima fused to two COX8 mitochondrial targeting signals in tandem (mt-mKeima). For transfections into the landing pad, the Bxb1 recombinase was expressed from pCAG-NLS-HABxb1 (Addgene #51271).

The Parkin variant library was based on the *PRKN* VAMP-seq library previously described^29^. From this library the mCherry was replaced with mt-mKeima. Mt-mKeima was amplified with eight PCR reactions with 10 ng of the p3024 plasmid (made by Genscript containing the *PRKN* WT cDNA) each, by using primers VV232 and VV233. The PCR program used was: 98 °C for 30 s, 12 cycles of 98 °C for 5 s, 66 °C for 30 s, 72 °C for 40 s and a final step at 72 °C for 5 min. The *PRKN* VAMP-seq library was amplified with primers flanking the mCherry sequence to produce linear vectors without mCherry, where mt-mKeima could be inserted. Eight reactions were performed with the primers VV230 and VV231 and the program was: 98 °C for 30 s, 10 cycles of 98 °C for 5 s, 66 °C for 30 s, 72 °C for 5 min and a final step at 72 °C for 5 min. For all PCRs the Q5 master mix (New England Biolabs) was used, following the protocol provided by the manufacturer.

The eight PCR reactions of mt-mKeima were mixed in one tube and were purified using Zymo Clean and Concentrator-5 kit (Zymo Research). The product was then eluted in 20 μL of nuclease-free water (Thermo Fisher Scientific). Four reactions of the vector PCR product were mixed into two tubes and they were purified by the Zymo Clean and Concentrator-5 kit (Zymo Research) and eluted in 16 μL nuclease-free water (Thermo Fisher Scientific). To remove any remaining template from the vector PCR product, it was treated with 2 μL DpnI (Thermo Fisher Scientific) for 2 h at 37 °C. The digested vector PCR product and the purified mt-mKeima product were separated on an agarose gel (1% for vector PCR and 2% for mt-mKeima). DNA was stained with 1x SYBR Safe (Thermo Fisher Scientific). The DNA bands of the correct size were excised from the agarose gels and purified using the GeneJET Gel Extraction Kit (Thermo Scientific). The concentration of the purified DNA was determined with the Qubit 1X dsDNA High Sensitivity (HS) Kit (Thermo Fisher Scientific).

The flanking sequences of the vector and of mt-mKeima products were amplified, so that they are compatible for Gibson assembly. The vector was mixed with mt-mKeima in a 1:4 molar ratio in five tubes and five Gibson assembly reactions were performed, using the Gibson assembly Master Mix (New England Biolabs). The five reactions were mixed and purified with the Zymo Clean and Concentrator-5 kit (Zymo Research), and they were eluted in 30 μL nuclease-free water (Thermo Fisher Scientific).

From the purified Gibson assembly reactions, 2 μL were electroporated into 50 μL NEB-10β *E. coli* cells (New England Biolabs) at 2 kV. The transformation was performed 11 times to ensure complete coverage of the library. A small portion of the transformed cells (1:110,000) were plated on an LB-agar plates with ampicillin (100 μg/mL) and left to grow at 37 °C overnight. The rest of the cells were inoculated into two flasks with 200 mL of LB media containing ampicillin (100 μg/mL) and the cultures were incubated, shaking (250 rpm) overnight at 37 °C. The next day the colonies growing on the LB-agar plate were counted and the liquid cultures were used to perform midi prep using the NucleoBond Xtra Midi kit (Macherey-Nagel). Based on the CFU number, the PRKN variant library complexity was covered 4,000-fold. The DNA yield and purity from the midiprep was measured by the NanoDrop spectrometer ND-1000.

### Cell culture and maintenance

We used HEK293T TetBxb1BPFiCasp9 Clone 12 cells with a genomically integrated lentiviral landing pad^33^. The cells were cultured in Dulbecco’s Modified Eagle High-Glucose medium (Sigma-Aldrich), supplemented with 10% (v/v) fetal bovine serum (FBS) (Sigma Aldrich), 5 mg/mL streptomycin (BioChemica), 5000 IU/mL penicillin (BioChemica), and 2 mM L-glutamine (Sigma Aldrich). For inducing expression of the landing pad system, cell cultures were further supplemented with 2 µg/mL doxycycline (Dox) (Sigma Aldrich). The cells were grown in a humidified incubator at 37 °C and 5% CO_2_. The cells were passaged every time they reached 70-80 % confluency and fresh Dox was added at each passage. Cells were kept in culture for a maximum of 20 passages. The cells were tested negative for mycoplasma. Cell line authentication was performed by checking for expression of BFP from the Tet-on promoter in non-recombinant cells.

### Transfections

Single Parkin variants, or the Parkin variant library, were transfected into HEK293T TetBxb1BPFiCasp9 Clone 12 cells and integrated in the genomic landing pad through Bxb1 specific recombination using FuGENE HD transfection reagent (Promega, E2312). The manufacturer’s protocol was followed, using a ratio of 1:14 of Bxb1 plasmid to Parkin plasmid, and a ratio of 1:4 of DNA mix to transfection reagent. After 48 hours, 10 nM AP1903 (MedChemExpress) was added to induce apoptosis in non-recombinant cells^33^. At the same time 2 μg/mL Dox was added to induce expression from the landing pad. After the selection, cells were kept in culture for 4 days with one passage at the 2-day mark. This allowed the cells to reach a steady-state expression level from the landing pad before further analysis.

### Fluorescence microscopy

After the successful integration of the Parkin variant, 8×10^4^ cells were seeded in 24-well plates (Costar) 3 days after the induction of expression from the landing pad. Cells were allowed to grow for an additional 48 hours before treatment with fresh DMEM containing 250 nM MitoTracker Red CMXRos (Invitrogen). After 30 minutes, mitophagy was induced by the addition of 2 µM antimycin A (Sigma-Aldrich) and 2 µM oligomycin (MedChemExpress) to the media or the respective volume of DMSO as control. Changes in GFP-Parkin localization were monitored using the Zeiss AX10 microscope with 10x objective. GFP and MitoTracker were excited by a 475 nm and 590 nm laser, respectively.

### Flow cytometry of single Parkin or ASPA variants

Four days after starting expression from the landing pad, the induction of mitophagy was achieved using 2 µM antimycin A (Sigma-Aldrich) and 2 µM oligomycin (MedChemExpress), or the corresponding volume of DMSO as a control, in fresh DMEM. When Bafilomycin A1 treatment was included, a 1 µM concentration was used and DMSO volume was increased accordingly. After treatment, the cells were washed with PBS, detached with 0.25 % trypsin (Gibco) and centrifuged at 300 g for 5 minutes. Finally, the cells were resuspended in PBS with 2 % FBS in round bottomed 96-well plates or filtered into flow cytometry tubes. Flow cytometry of single Parkin or ASPA variants in low throughput was carried out on the FACSCelesta or LSRFortessa X-20 flow cytometer (BD Biosciences), with a high throughput sampler for use with 96-well plates. The lasers and band filters were 488 nm and 530/30 for GFP, 405 nm and 450/40 for BFP, 561 nm and 610/20 nm for acidic mt-mKeima, and 405 nm and 610/20 for neutral mt-mKeima. The data were analyzed using the FlowJo v.10.10.0 software (BD Biosciences). Live and single cells were gated using forward scatter (FSC), side scatter and FSC pulse width. Then, BFP-negative and GFP-positive cells were gated for further analysis. If not stated otherwise, at least 7×10^3^ events were collected in the BFP-negative and GFP-positive population. The full gating strategy can be found in the supplementary material (**Supplementary Fig. 16**).

To display mitophagic activity, the acidic:neutral mt-mKeima mean of the mock treated cells (DMSO) was subtracted from the mean of the AO treated cells to remove background signal. Then, the corrected mitophagy signal for each variant was normalized to Parkin or ASPA WT activity within the respective replicate measurement.

### FACS

The Parkin library was transfected into the HEK293T TetBxb1BPFiCasp9 Clone 12 cells as described above. To induce mitophagy, the cells were treated for 4 hours with either 2 μM antimycin A and 2 µM oligomycin A (AO), or with the equivalent volume of DMSO as a mock treatment. The cells were then washed with PBS, detached with trypsin, resuspended in PBS with 2% FBS, and filtered through a 50 µm nylon mesh filter (BD Biosciences). Cell sorting was carried out on a BD FACSymphony S6 (BD Biosciences) equipped with a 70 μm nozzle. The lasers used were 488 nm (GFP), 405 nm (BFP), and 561 nm, along with 530/30 nm (GFP), 431/28 nm (BFP), 610/20 nm, and 605/40 nm band pass filters, and 505 nm (GFP), 410 nm (BFP), 600 nm, and 595 nm long pass filters. The ratio of acidic mt-mKeima to neutral mt-mKeima was used as a metric for mitophagy levels. Acidic mt-mKeima was detected using excitation at 561 nm and emitted light was collected after passing through a 600 nm long pass filter and a 610/620 nm band pass filter, while the neutral mt-mKeima was detected using excitation at 405 nm and emitted light was collected through a 595 nm long pass filter and a 605/40 nm band pass filter. Cells were gated for live cells and singlets, that were BFP negative and mt-mKeima positive. A full gating strategy can be found in the supplementary material (**Supplementary Fig. 17**). A histogram of the acidic:neutral mt-mKeima ratiometric parameter was determined on the BD FACSDiva software and gates were set to sort the entire library into four bins based on the ratios. The bins were of unequal sizes with the first bin covering 15% of the cells with the lowest ratio, the second bin covering 30% of the total population, the third bin covering 35% of the total population, and the fourth bin covering the 20% with the highest ratio. Proportionally to the bin percentages, 2×10^6^ cells were sorted from the lowest bin, 4×10^6^ cells from the second lowest, 4.67×10^6^ cells from the second highest, and 2.67×10^6^ cells from the highest bin. Throughout the entire protocol a minimum of 1×10 cells was maintained at each step to ensure at least 100-fold library coverage. To have enough DNA for sequencing in the following steps, the number of cells sorted into the 15% bin was increased to 2×10^6^ and the number of cells in the other bins scaled up accordingly to match the percentages. Cell sorting was repeated for a total of three independent replicates both with AO and DMSO treatment (separate library transfections, a total of six sorts). After sorting, the samples were pelleted by centrifugation at 300 g for 5 minutes, the supernatant removed, and the pellets were placed for storage at -80 °C until further processing. Further analysis of FACS data was performed using the FlowJo v.10.10.0 software (BD Biosciences).

### Genomic DNA extraction

Genomic DNA was extracted from sorted cells using the DNeasy blood & tissue Kit (Qiagen). The manufacture’s protocol, *Purification of Total DNA from Animal Blood or Cells (Spin-Column Protocol)*, was followed with the following exceptions: Addition of RNase was reduced to 0.2 mg per sample. The cell lysing incubation was extended from 10 to 30 minutes with shaking. PCR grade water was used instead of elution buffer. Samples from the 15% and 20% FACS bins were eluted in 100 µL, and the other samples in 150 µL. The eluate was passed through the column twice to increase the yield.

### Genomic DNA amplification, purification, and quantification

Genomic DNA was amplified with PCR. The number of reactions was kept proportional to the bin percentage for each sample, with the samples from the 15% bins having 3 reactions, samples from the 30% bins having 6 reactions, samples from the 35% bins having 7 reactions, and samples from the 20% bins having 4 reactions. For each reaction 1.5 µg of DNA was mixed with 25 µL 2x Q5 High-Fidelity Mastermix (New England BioLabs), 0.5 µM of primers LC1020 and LC1031 (**Supplementary Table 5**), and PCR grade water to a final volume of 50 µL. The initial denaturation was carried out at 98 °C for 30 s, followed by 7 cycles of denaturation at 98 °C for 10 s, annealing at 60 °C for 20 s, and extension at 72 °C for 15 s. A final extension was carried out at 72 °C for 2 minutes before holding at 4 °C.

After the PCR amplification, the reactions for each sample were pooled together in one 1.5 mL Eppendorf tube, and AMPure XP Beads (Beckman Coulter) were used for PCR product purification. A 1:1 ratio of bead volume to PCR product volume was used, 4x the volume of the pooled PCR product of fresh 70% ethanol was used for washing, and DNA was eluted in 21 µL of PCR grade water.

To shorten the amplicon and add P5 and P7 cluster-generating sequences and indexes for Illumina sequencing, a qPCR reaction was performed with the bead purified PCR samples. For each reaction 8 µL of the purified DNA were mixed with 25 µL 2x Q5 High-Fidelity Mastermix (New England BioLabs), 10x SYBR green (Invitrogen), 0.5 µM of RCR2_Fw1 I5 indexing primer, 0.5 µM of a JS_R I7 primer with a unique index to each sample (**Supplementary Table 5**), and 9.5 µL PCR grade water. The initial denaturation was carried out at 98 °C for 30 s. Then the program was set to cycle through the following steps: denaturation at 98 °C for 10 s, annealing at 63 °C for 20 s, and extension at 72 °C for 15 s. The reaction was monitored in real time and stopped after 16 cycles, when the amplification curve was in early exponential phase. Finally, the amplicons were separated on a 2% agarose gel with 1x SYBR Safe (Invitrogen) and extracted using the GeneJet gel extraction kit (Thermo Scientific, K0692) according to the manufacturer’s protocol. 30 µL PCR grade water were used for the final elution. The extracted amplicons were quantified with the Qubit 2.0 fluorometer, using the Qubit dsDNA HS Assay Kit (Invitrogen).

### Sequencing

For sequencing, the amplicons were pooled keeping the molarities proportional to the percentage of the bin they originated from. The pooled library was then denatured, diluted and mixed with PhiX Control v3 according to the manufacturer’s protocol (Illumina). The library sequencing was performed using a NextSeq550 Illumina sequencer and a NextSeq 500/550 High Output v2.5 75 cycle kit. Custom sequencing primers were used: LC1040 for Read 1, LC1041 for Read 2 (paired-end), LC1042 for Index 1 and ASPA_PARK2_index2_re for Index 2 (**Supplementary Table 5**).

### Data analyses

Sequencing reads of barcodes were counted and mapped as in Clausen *et al*. 2024^29^. Briefly, adapter sequences were removed using cutadapt^89^ and pair-end reads were joined using fastq-join from EA utils^90^. Only exact barcode matches were counted and merged per amino acid variant. We obtained a synonymous count per amino acid position by merging all counts that map to a silent single-nucleotide substitution in each codon. For each biological replicate we calculated average bin scores using a pseudo count of one and weighted each gate according to population (**Supplementary Fig. 2**). Average bin scores based on less than 50 reads were discarded. A score per amino acid variant for AO and DMSO treated cells was obtained by averaging replicates requiring at least two replicate measurements.

To obtain an AO-induced activity score, we subtract the signal from the DMSO treated cells from the signal of the AO treated cells. However, these scores are not necessarily on the same scale since the FACS gating and fluorescence distributions are different. To inform a scaling we assume that nonsense variants are on average as inactive as the catalytic-site control variant C431A. This resulted in a DMSO scaling of 1.77 (**Supplementary Fig. 6A-D**). After subtracting the scaled DMSO score from the AO score, a final activity score was obtained by min-max normalization to WT and C431A (by construction equal to the average score of nonsense variants).

In an alternative approach, both AO and DMSO average bin scores are transformed to match their respective FACS log10(mt-mKeima acidic:neutral) ratio distributions using a non-linear and rank preserving scaling method described previously^49^. On this fluorescence scale, scores should be comparable allowing direct subtraction of the (parkin-independent) DMSO mitophagy from the AO induced mitophagy (**Supplementary Fig. 6D-F**). These fluorescence-transformed scores support the assumption that nonsense variants are on average inactive and are available in the supporting material (**Supplementary Data File**).

### Predicted properties and clinical data

Predictions of structural stability using Rosetta and evolutionary conservation using GEMME are from Clausen *et al*^29^. ESM-IF and ESM1b are from Cagiada *et al*^31^. AlphaMissense^58^ and popEVE^57^ scores were downloaded via links in the original descriptions.

Allele frequencies from GnomAD v4.1.0^61^ are based on 721 missense variants from transcript ENST00000366898.6. For ten amino acid variants with more nucleotide substitutions registered, we obtain a merged frequency by adding the allele counts and dividing by the allele number associated with the highest allele count, assuming these are the approximately same populations and taking the conservative choice. This resulted in allele frequencies for 711 amino acid variants.

Clinical data from ClinVar were obtained from Cagiada *et al*. 2024^91^ and combined with annotations from MDSgene^59^. In particular, if ClinVar annotated a variant as (likely) pathogenic or (likely) benign, then the ClinVar annotation was used. If ClinVar had a VUS, conflict or no annotation for a variant and MDSgene had an annotation for that variant, then the MDSgene annotation was used.

C-degron effects were predicted using the peptide abundance predictor (PAP) using the cnn2w1 model set at a tile size of 30, including the calculation of C-degron scores, without using the full-coverage or the saturation mutagenesis methods^49^.

## Acknowledgements

We acknowledge the use of the FACS, sequencing and computing core facilities at the Department of Biology, University of Copenhagen. We thank Anne-Marie Lauridsen and Celeste M. Hackney for technical assistance. We thank Prof. Morten Meyer for helpful discussions and Prof. Dr. Katja Lohmann and MDSGene for providing helpful information about *PRKN* variants. Fig. 1A was made using BioRender.com.

## Abbreviations

ACT: activation element
AO: antimycin and oligomycin
AM: AlphaMissense
ARPD: autosomal-recessive Parkinson’s disease
AUC: area under the curve
Baf: Bafilomycin
DMEM: Dulbecco’s modified Eagle medium
Dox: doxycycline
DUBTAC: deubiquitylating targeted chimeras
FACS: fluorescence-activated cell sorting
FBS: fetal bovine serum
IBR: in-between RING
IQR: interquartile range
IRES: internal ribosomal entry site
MAGIC: mitochondria as guardian in cytosol
MAVE: multiplexed assay of variant effects
MED: median
OMM: outer mitochondrial membrane
PAP: peptide abundance predictor
PD: Parkinson’s disease
REP: Repressor element
ROC: Receiver Operating Characteristic
SD: standard deviation
pUb: phospho-ubiquitin
pUBL: phospho-UBL
Ub: ubiquitin
UBL: ubiquitin-like
VAMP-seq: variant abundance by massively parallel sequencing
VUS: variant of unknown significance
WT: wild-type.

## Ethics approval and consent to participate

Not applicable.

## Consent for publication

Not applicable.

## Supplemental material

This article includes the following supplemental information.

- Supplementary figures and tables (SupplementaryInformation.pdf).
- Supplementary data file (SupplementaryDataFile.xlsx).

## Data availability

All data and software generated for this article are available on GitHub (https://github.com/KULL-Centre/_2026_Sigmarsdottir_parkin_activity). The sequencing data are available at the NCBI sequence read archive (SRA) BioProject PRJNA1359748 and on ERDA (https://sid.erda.dk/sharelink/ff6CDV5565). The mutational map is available at MaveDB (https://www.mavedb.org/).

## Author contributions

E.S.S., V.V., K.E.J., A.B. and I.K.H. performed the experiments. E.S.S., V.V., K.E.J., I.K.H., K.L.-L., and R.H.-P. analyzed the data. V.V., K.L.-L. and R.H.-P. conceived the study, E.S.S., V.V. and R.H.-P. wrote the paper.

