## Supplementary figures and tables for "Comprehensive Variant Effect Map of Parkin-Mediated Mitophagy in Parkinson’s Disease"

### *Supplementary information*


**Suppl. Fig. 1** – *Localization of WT Parkin and the C431A variant.* p.2
**Suppl. Fig. 2** – *FACS of DMSO treated cells.* p.3
**Suppl. Fig. 3** – *DMSO affects mitophagy.* p.4
**Suppl. Fig. 4** – *Variant effect map of Parkin activity under DMSO treatment.* p.5
**Suppl. Fig. 5** – *Variant effect map of Parkin activity under AO treatment.* p.6
**Suppl. Fig. 6** – *Combining AO and DMSO screens to AO-induced activity scores.* p.7
**Suppl. Fig. 7** – *Reproducibility of average bin scores between replicates.* p.8
**Suppl. Fig. 8** – *Data filtering based on the standard deviation.* p.9
**Suppl. Fig. 9** – *Mitophagy induced by low abundance Parkin variants.* p.10
**Suppl. Fig. 10** – *Correlation with C-degron potency of nonsense variants in IDR.* p.11
**Suppl. Fig. 11** – *Mitophagy is induced by misfolded protein variants.* p.12
**Suppl. Fig. 12** – *Variants with low abundance and high activity are poorly predicted.* p.13
**Suppl. Fig. 13** – *Activity and abundance scores of variants based on domain positioning.* p.14
**Suppl. Fig. 14** – *Comparisons with variant effect predictors.* p.15
**Suppl. Fig. 15** – *Activity and Alpha Missense predictions of pathogenic variants.* p.16
**Suppl. Fig. 16** – *Gating strategy used for flow cytometry.* p.17
**Suppl. Fig. 17** – *Full gating strategy for FACS of the Parkin variant library* p.18

**Suppl. Table 1** – *Variants selected for low throughput validation.* p.19
**Suppl. Table 2** – *Notable positions in Parkin.* p.20
**Suppl. Table 3** – *Positions without known mechanisms.* p.22
**Suppl. Table 4** – *Notable positions where substitutions cause minor effects.* p.23
**Suppl. Table 5** – *Primers used in this study.* p.24

**Suppl. References** p.25

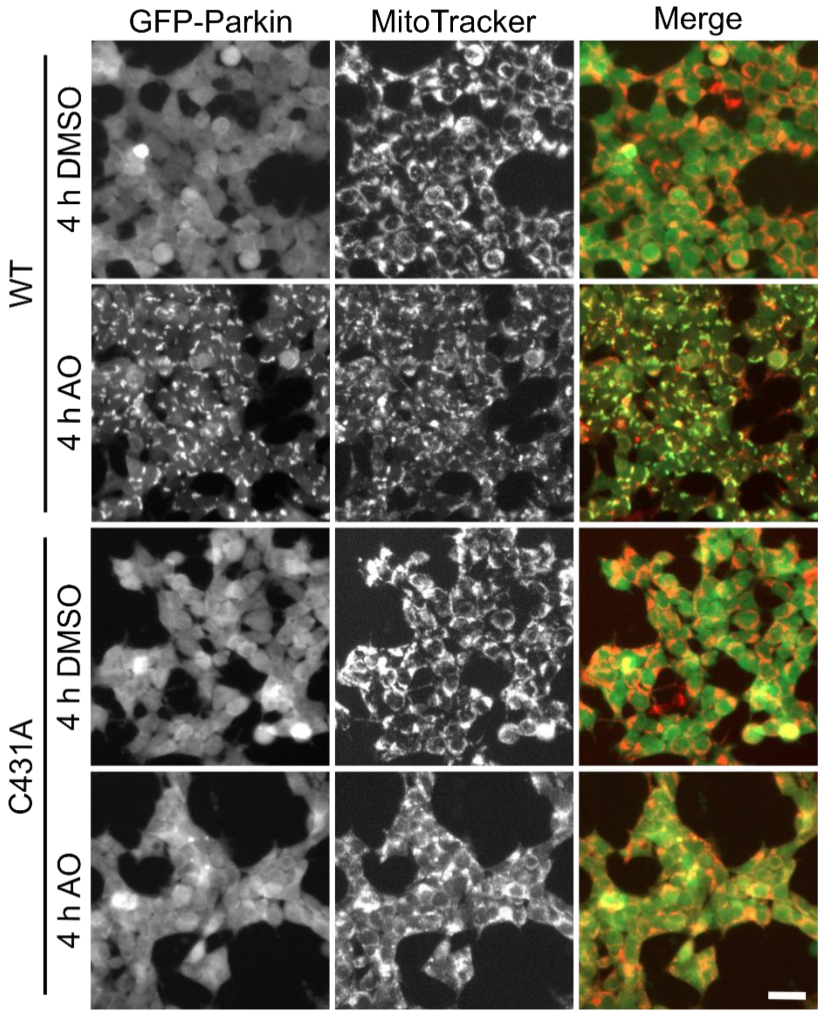


**Supplementary Figure 1.** *Localization of WT Parkin and the C431A variant.* HEK293T landing pad cells stably expressing either wild-type (WT) or catalytically dead C431A GFP-fused Parkin were pre-treated with 250 nM MitoTracker for 30 minutes. Afterwards, the cells were incubated with 2 μM antimycin and oligomycin (AO) or the corresponding volume of DMSO for 4 hours. GFP-Parkin and mitochondria localization was monitored by live cell fluorescence microscopy. Scale bar, 30 μm.

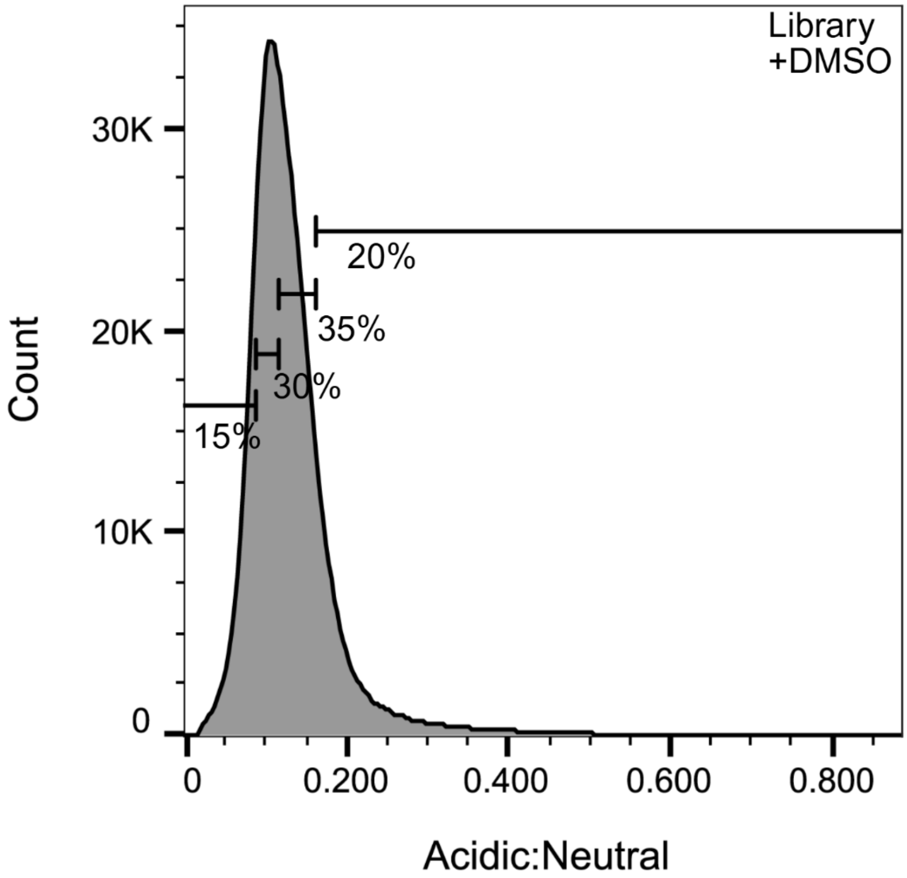


**Supplementary Figure 2.** *FACS of DMSO treated cells.* The DMSO-treated Parkin variant library expressing cells were, like the AO treated population, sorted into four bins based on their acidic:neutral mtKeima ratio. The range was much narrower for cells treated with DMSO compared to the AO treated cells (**Fig. 1D**), as mitophagy was not induced, but the percentages of the population in each bin were kept consistent between the two conditions. The lowest bin was set to cover 15%, the second to cover 30%, the third to cover 35%, and the highest to cover 20% of the whole population.

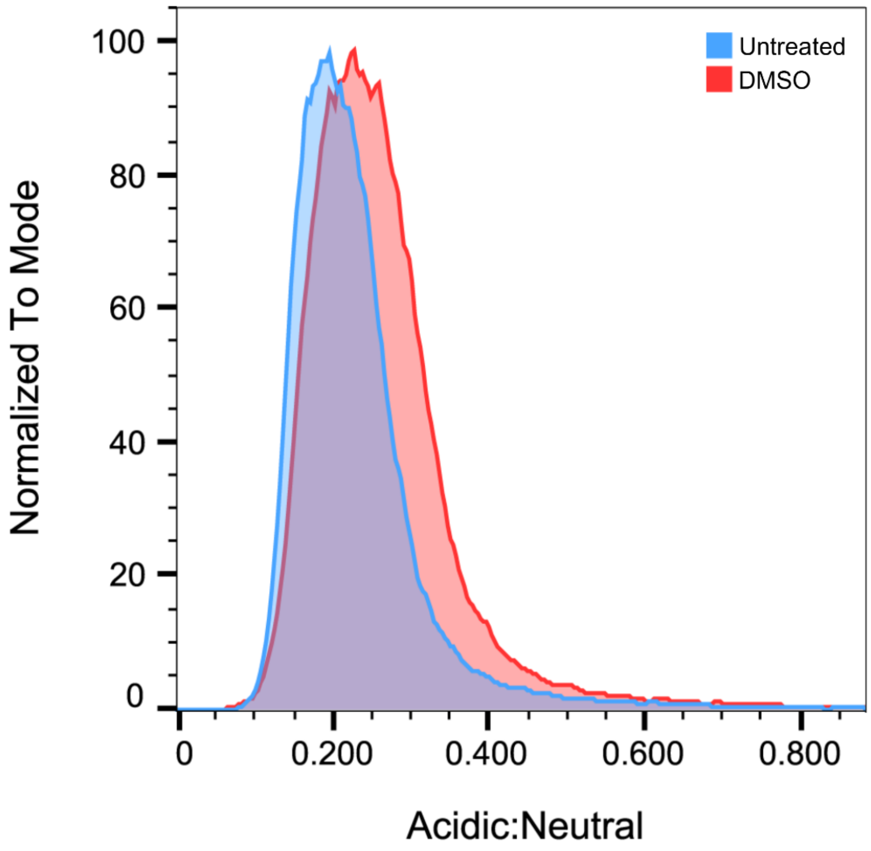


**Supplementary Figure 3.** *DMSO affects mitophagy.* Cells expressing the Parkin variant library were treated with either DMSO (red, n=87,548) or left untreated (blue, n=88,705). The effect of DMSO treatment on mitophagy was assessed by measuring the acidic:neutral mtKeima ratio by flow cytometry. The profiles show that DMSO treatment induced a low level of mitophagy. This background mitophagy was accounted for by subtracting the mitophagy level of the DMSO treated samples from the AO treated samples during data analysis.

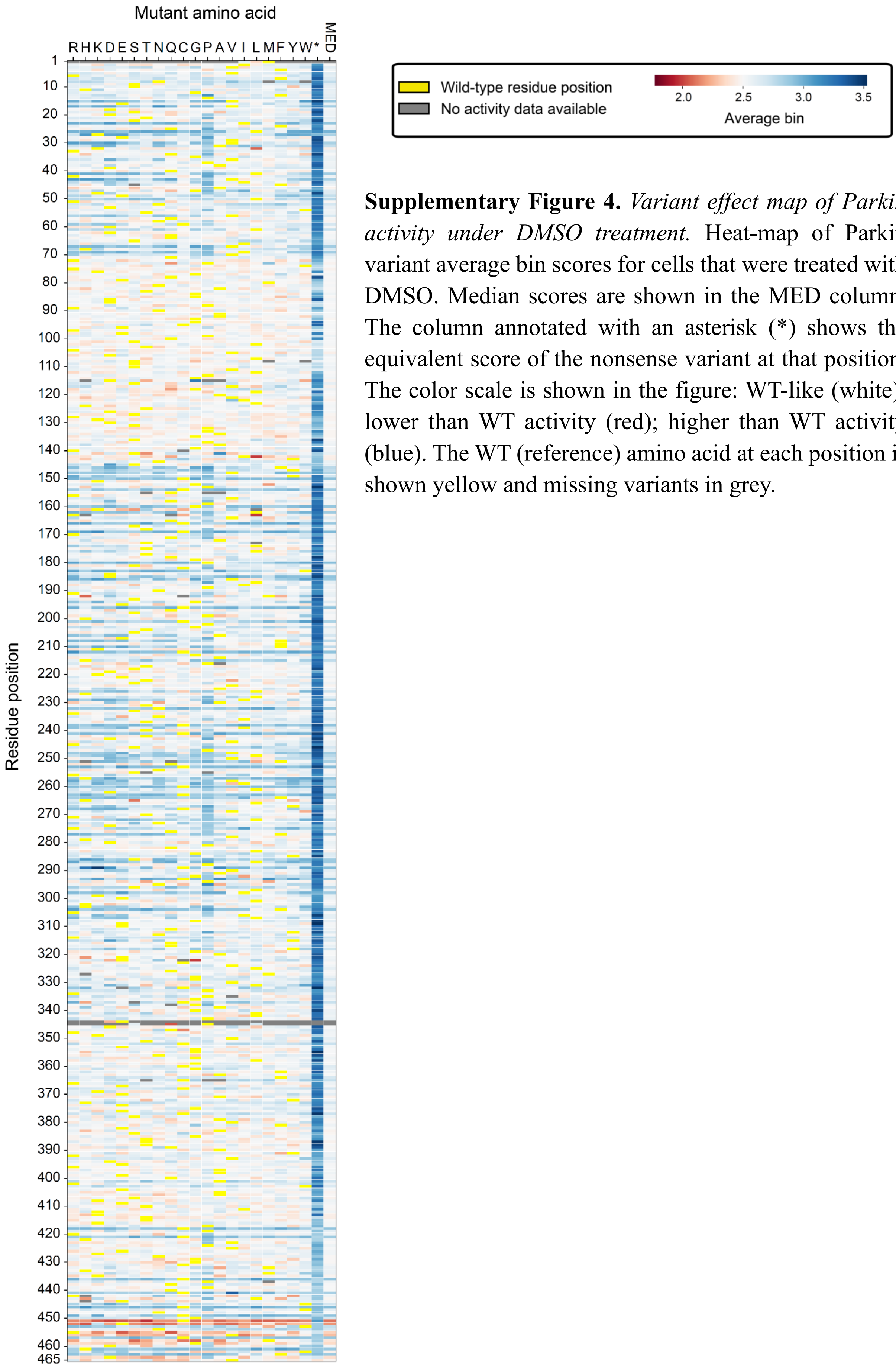


**Supplementary Figure 4.** *Variant effect map of Parkin activity under DMSO treatment.* Heat-map of Parkin variant average bin scores for cells that were treated with DMSO. Median scores are shown in the MED column. The column annotated with an asterisk (*) shows the equivalent score of the nonsense variant at that position. The color scale is shown in the figure: WT-like (white); lower than WT activity (red); higher than WT activity (blue). The WT (reference) amino acid at each position is shown yellow and missing variants in grey.

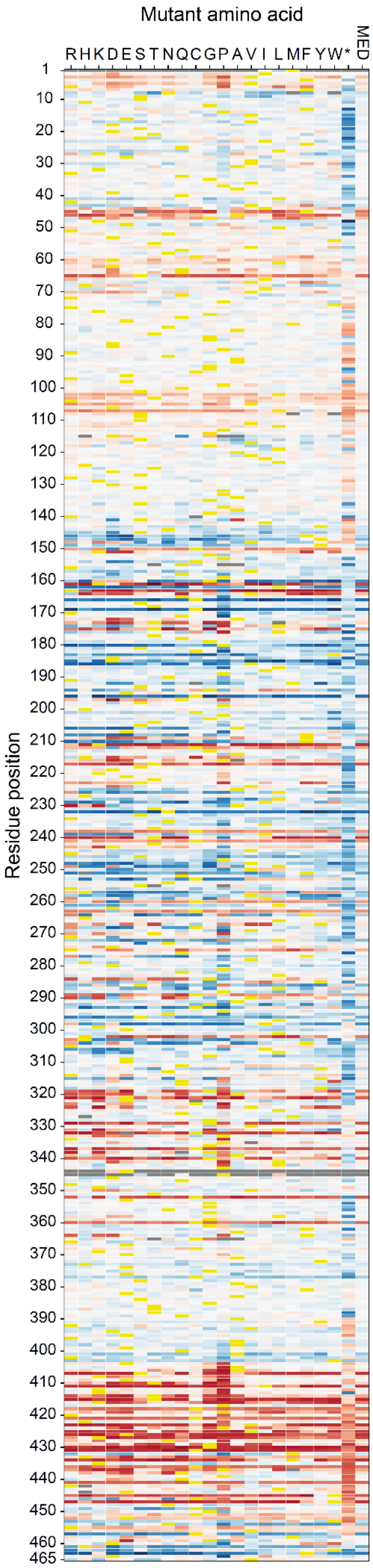


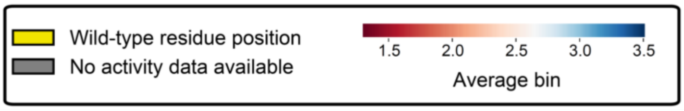


**Supplementary Figure 5.** *Variant effect map of Parkin activity under AO treatment.* Heat-map of Parkin variant average bin scores for cells that were treated with 2 μM of antimycin and 2 μM of oligomycin (AO). Median scores are shown in the MED column. The column annotated with an asterisk (*) shows the equivalent score of the nonsense variant at that position. The color scale is shown in the figure: WT-like (white); lower than WT activity (red); higher than WT activity (blue). The WT (reference) amino acid is shown in yellow and missing variants in grey.

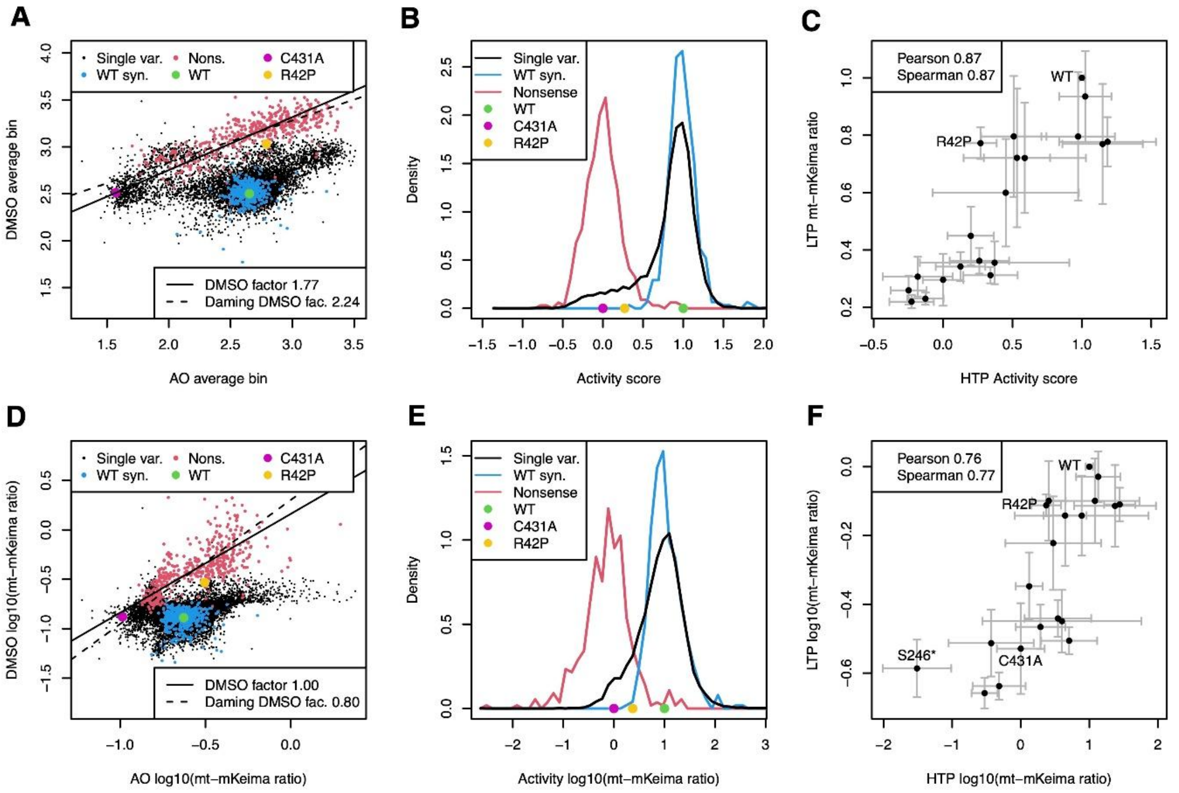


**Supplementary Figure 6.** *Combining AO and DMSO screens to AO-induced activity scores.* The majority of nonsense variants were screened to be active or even hyperactive both with AO and DMSO treatment (A), so we subtracted the DMSO signal from the AO signal to obtain an AO-induced activity score. Two strategies are shown: A linear scaling of average bin scores assuming nonsense variants are inactive like C431A (A-C) or a non-linear transforming of both scores to a common scale of fluorescence ratios (D-F). The former was selected for the final scores. Scatter plots of the individual screens show a linear relation for nonsense variants (red points, Pearson 0.76 and 0.70 for panels A and D, respectively). For average bin scores a DMSO factor of 1.77 results in a mean score of nonsense variants that equals C431A (A) and for transformed fluorescence ratios, a scaling of one is assumed (panel D). Both scalings are similar to slopes obtained by Daming regression (line fit that assumes the same uncertainty in x and y). Score distributions (B,E) and low throughput (LTP) validations (C,F) are similar for the two methods with a slightly better separation between nonsense and synonymous WT, and LTP correlation for the average bin scores scaled by 1.77. The fluorescence transformed scores (D-F) confirm that the nonsense variants are indeed as inactive as C431A and thus support the assumption resulting in the scaling factor of 1.77 used in the final activity scores.

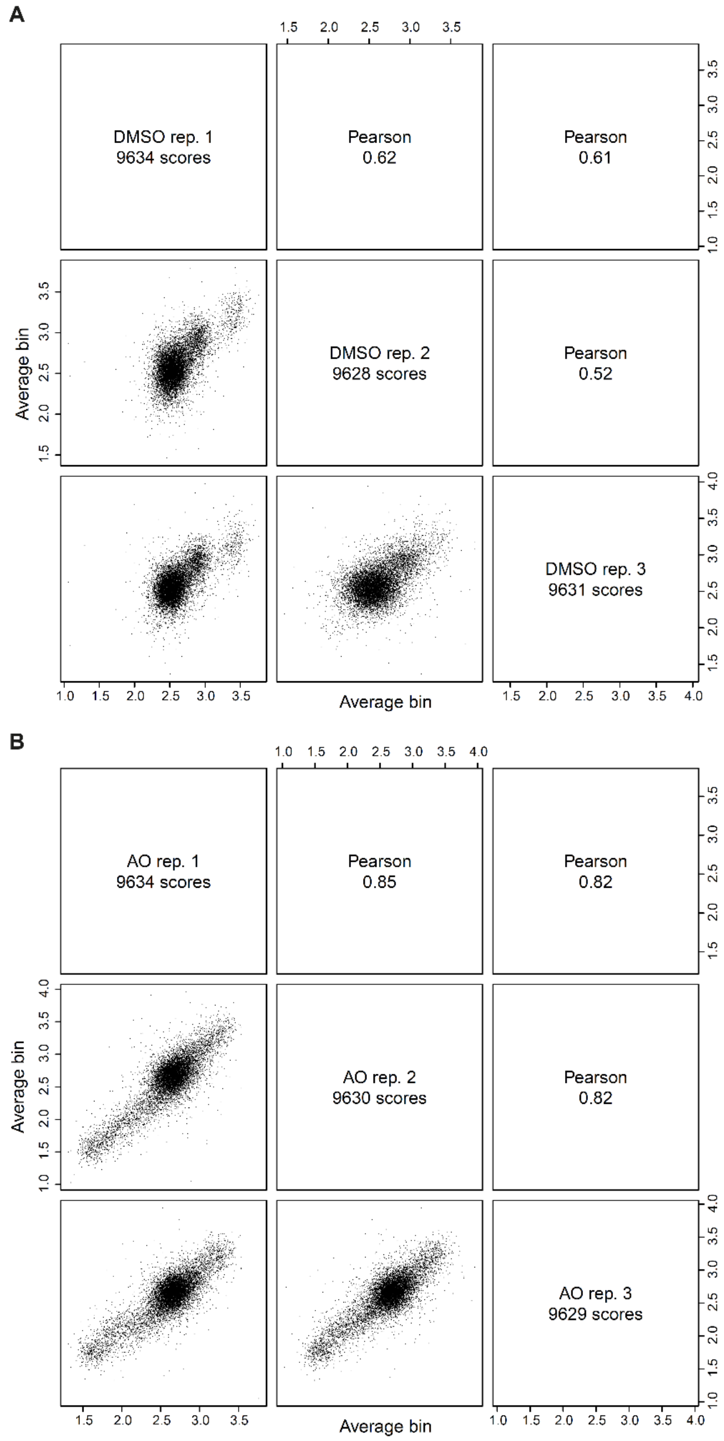


**Supplementary Figure 7.** *Reproducibility of average bin scores between replicates.* Pairwise correlations of the three replicates performed for Parkin activity under (A) DMSO or (B) AO (2 μM of antimycin and 2 μM of oligomycin) treatments. Pearson's r is indicated in the plot. Scores range from 1 to 4, with 1 indicating low Parkin activity and 4 indicating high Parkin activity.

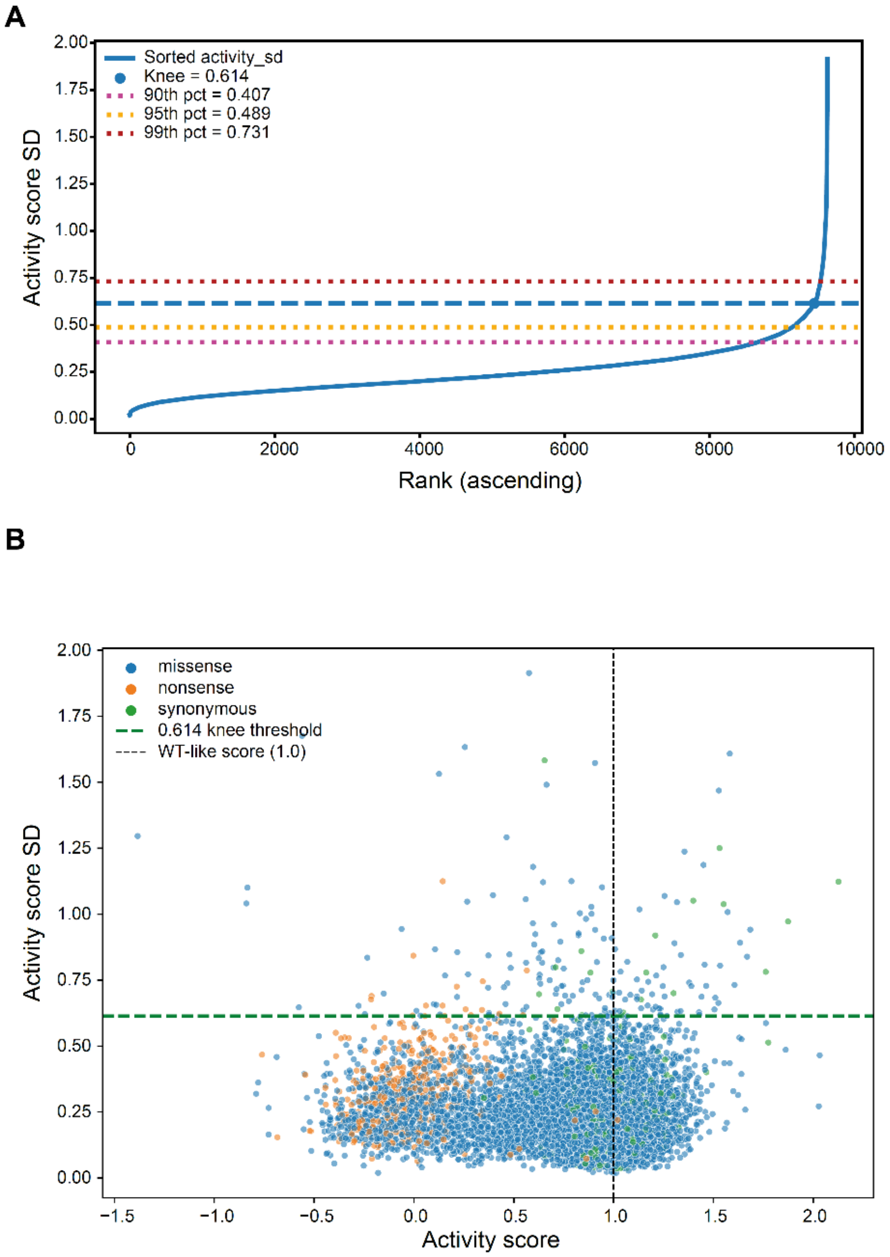


**Supplementary Figure 8.** *Data filtering based on the standard deviation.* (A) Data were filtered based on a standard deviation (SD) threshold that was determined by calculating the "knee" of the ranked distribution of SDs from all the datapoints. The "knee" value, and the 90$^{th}$, 95$^{th}$, and 99$^{th}$ percentiles are indicated for context. 9447/9632 (98.08%) datapoints pass the filter. (B) Scatterplot of activity score SD against activity score. Missense (blue), nonsense (orange) and synonymous (green) variants are highlighted. The cutoff threshold (green line) and the WT activity score (black line) are marked in the plot.

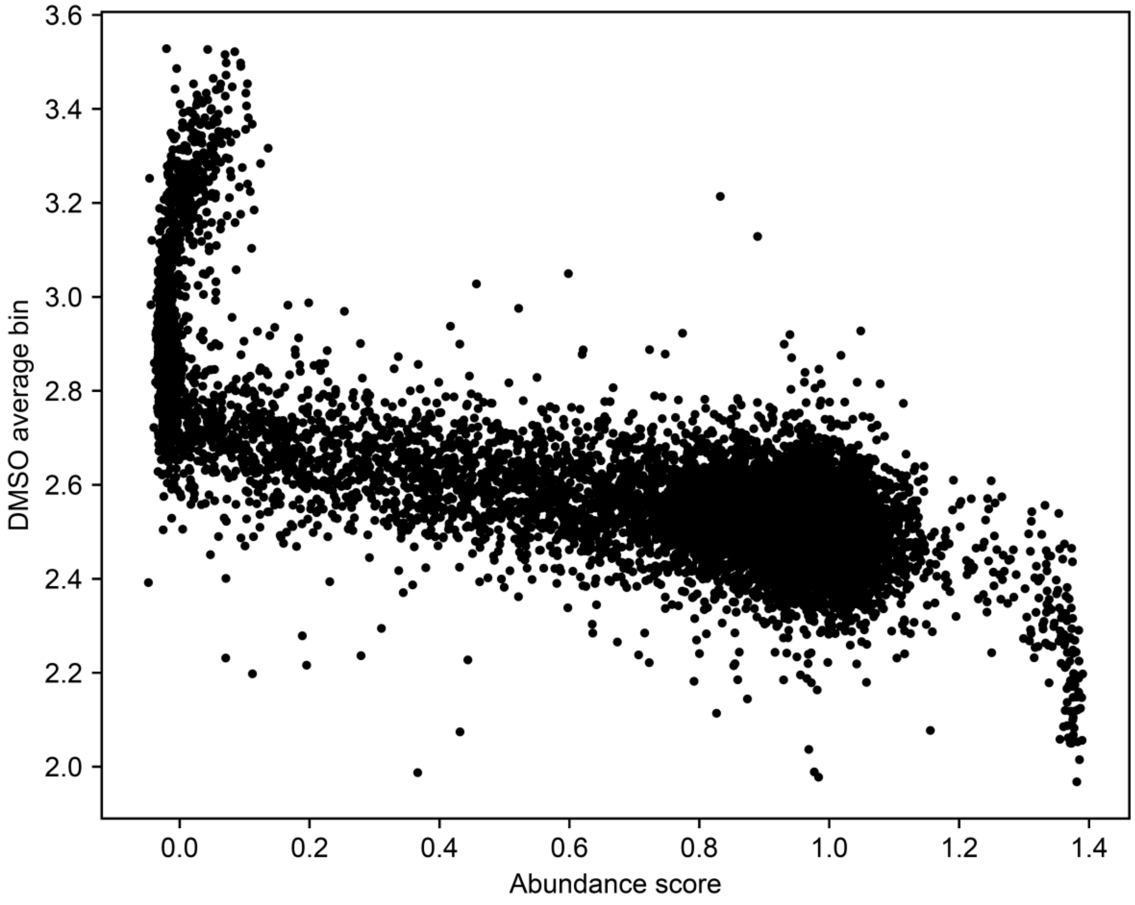


**Supplementary Figure 9.** *Mitophagy induced by low abundance Parkin variants.* Correlation between mitophagic activity of the variants in the library with DMSO treatment and their abundance[1]. Reduced abundance correlates with increased mitophagy with DMSO treatment (no AO induction).

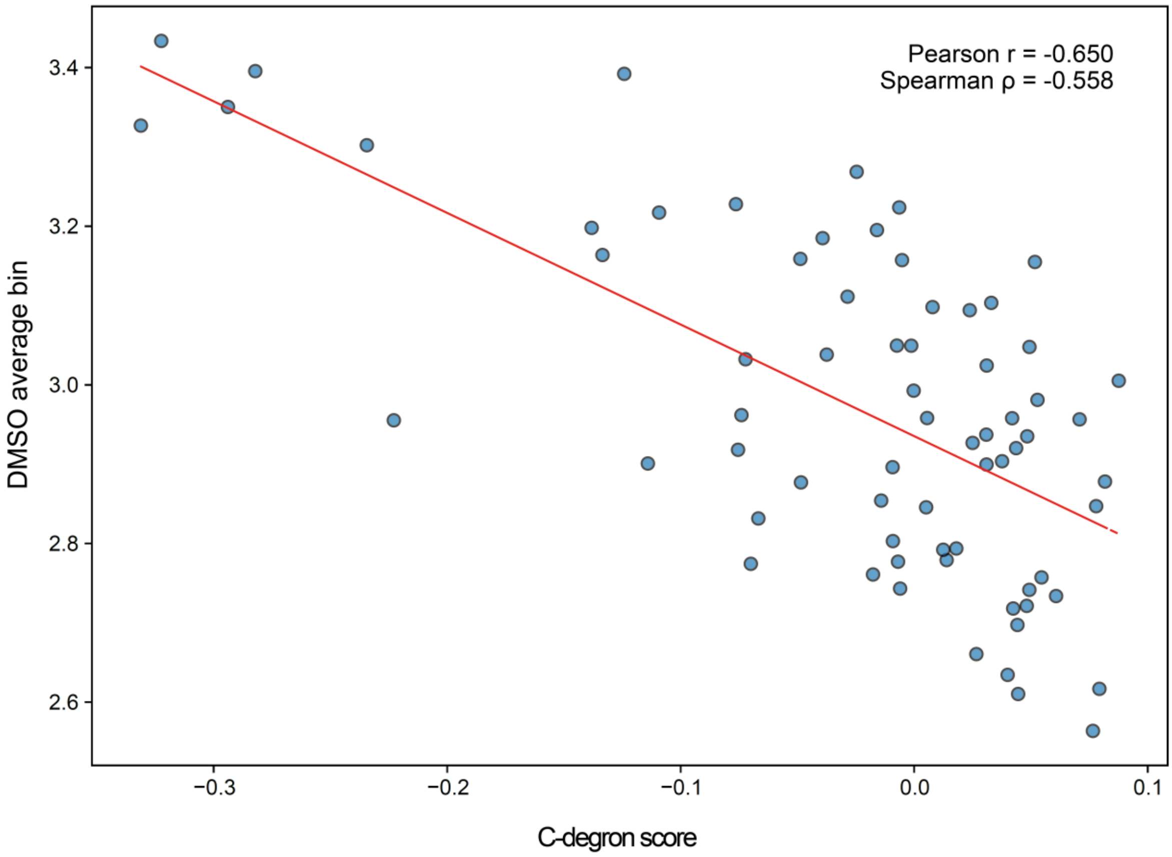


**Supplementary Figure 10.** *Correlation with C-degron potency of nonsense variants in IDR.* Scatterplot showing the correlation between the average bin scores under DMSO treatment and the C-degron score of Parkin nonsense variants within the disordered region of Parkin that links UBL and RING0 (positions 77-145). Low C-degron scores indicate high C-degron potency.

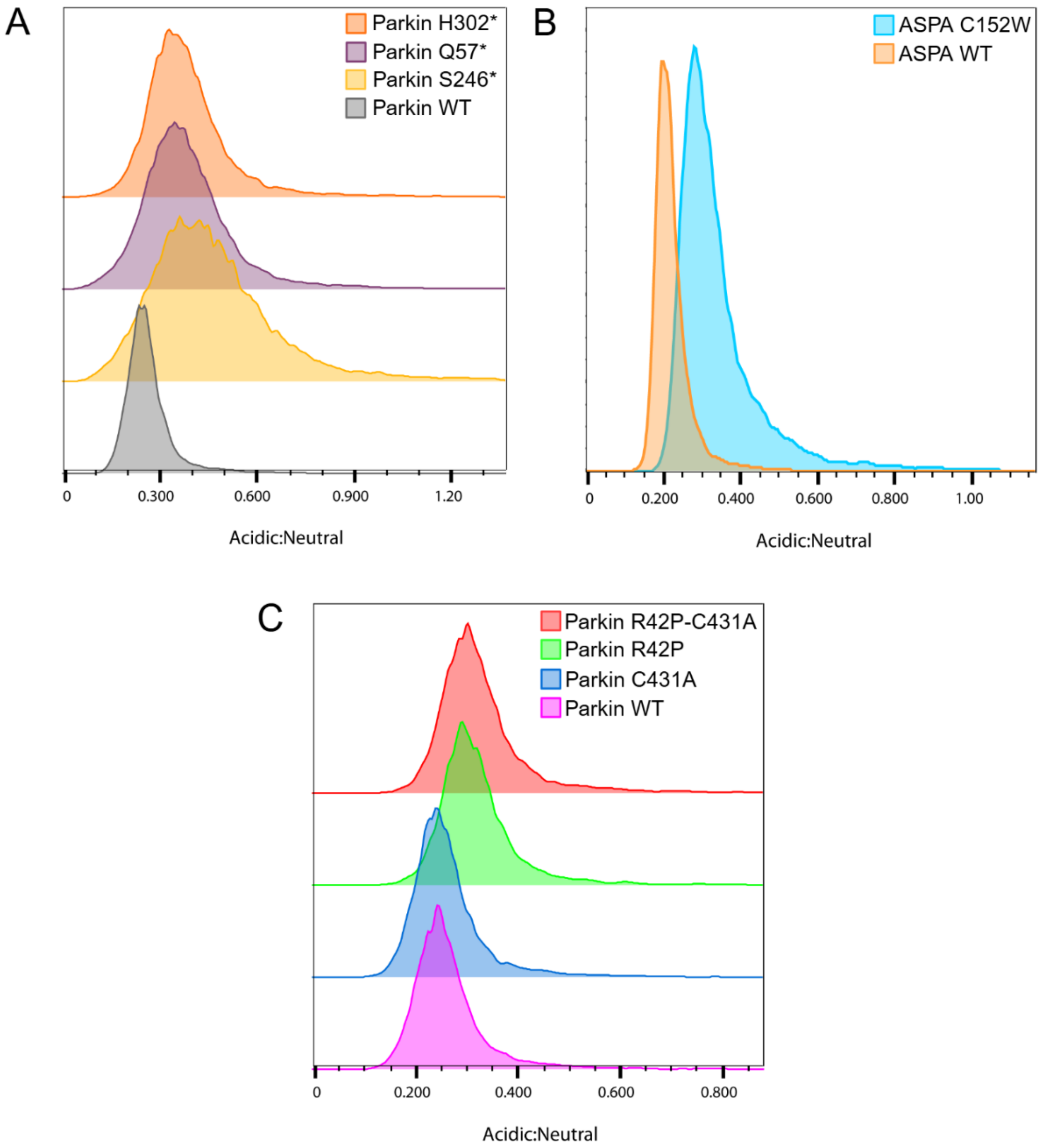


**Supplementary Figure 11.** *Mitophagy is induced by misfolded protein variants.* (A) Mitophagy levels in cells expressing Parkin nonsense variants (n>15,000) without mitophagy induction (DMSO mock treatment). Nonsense variants have higher background mitophagy compared to WT Parkin (grey, n=10,318). (B) Mitophagy levels in cells expressing either WT ASPA (orange, n=10,490) or the C152W ASPA variant (cyan, n=12,782). The C152W variant has heightened levels of mitophagy compared to ASPA WT, without AO induction of mitophagy (DMSO mock treatment). (C) Mitophagy levels of DMSO mock treated cells expressing Parkin R42P-C431A double mutant (red, n=11,996), Parkin R42P (green, n=11,825), Parkin C431A (blue, n=12,149), or Parkin WT (purple, n=11,378). Both the misfolded R42P variant and the R42P-C431A double mutant show elevated background mitophagy compared to WT.

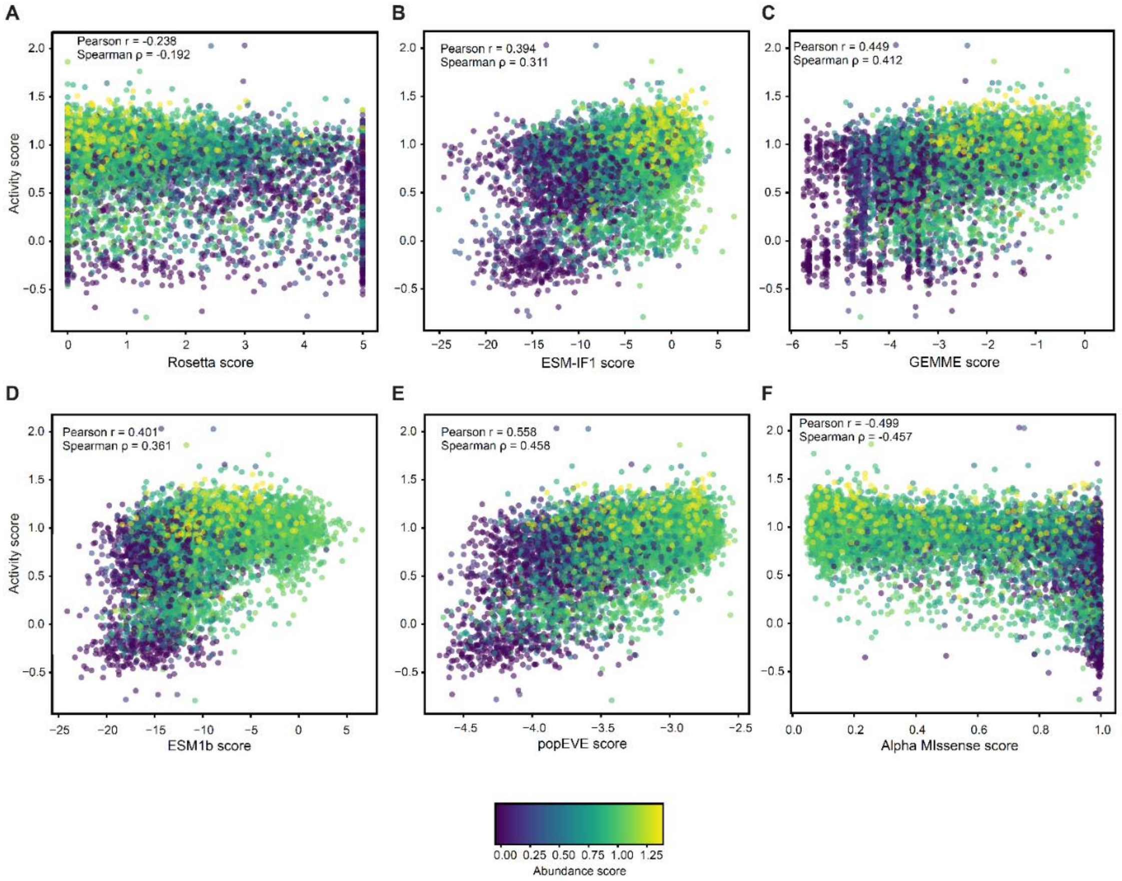


**Supplementary Figure 12.** *Variants with low abundance and high activity are poorly predicted.* (A-F) Scatterplots correlating Parkin activity scores with variant effect predictions based on (A) Rosetta, (B) ESM-IF1, (C) GEMME, (D) ESM1b, (E) popEVE, and (F) Alpha Missense. The data points are colored based on their abundance score from Clausen et al. (2024)[1]. The color scale is shown at the bottom.

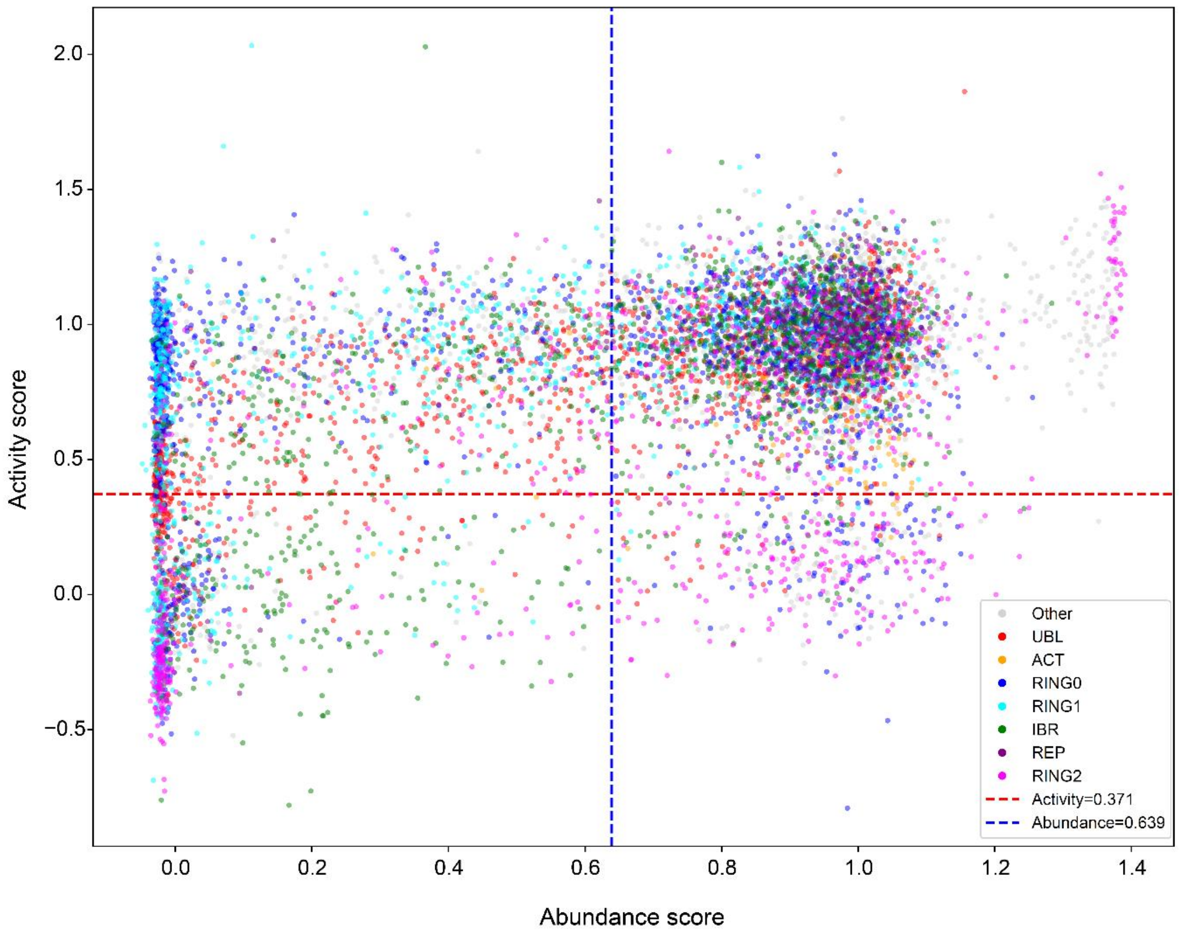


**Supplementary Figure 13.** *Activity and abundance scores of variants based on domain positioning.* Scatterplot showing the correlation of activity with abundance scores[1] of Parkin variants. Datapoints are colored based on the Parkin domain, at which they are located (UBL: red, ACT: orange, RING0: blue, RING1: cyan, IBR: green, REP: purple, RING2: magenta). The red and blue dotted lines show the thresholds that best separate the pathogenic/likely pathogenic from the benign/likely benign variants, according to Youden's J statistic.

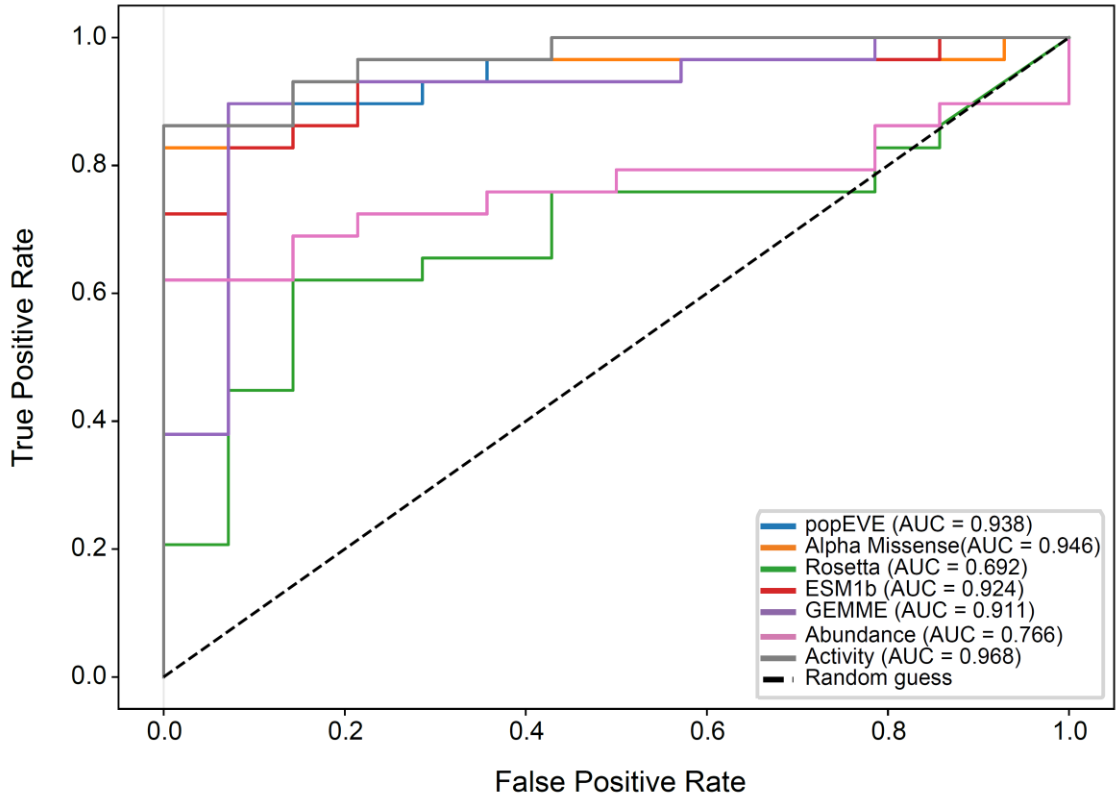


**Supplementary Figure 14.** *Comparisons with variant effect predictors*. ROC curves for activity score (grey), abundance score (magenta), GEMME score (purple), ESM1b score (red), Rosetta score (green), Alpha Missense score (orange) and popEVE score (blue). The AUC is reported in the figure legend.

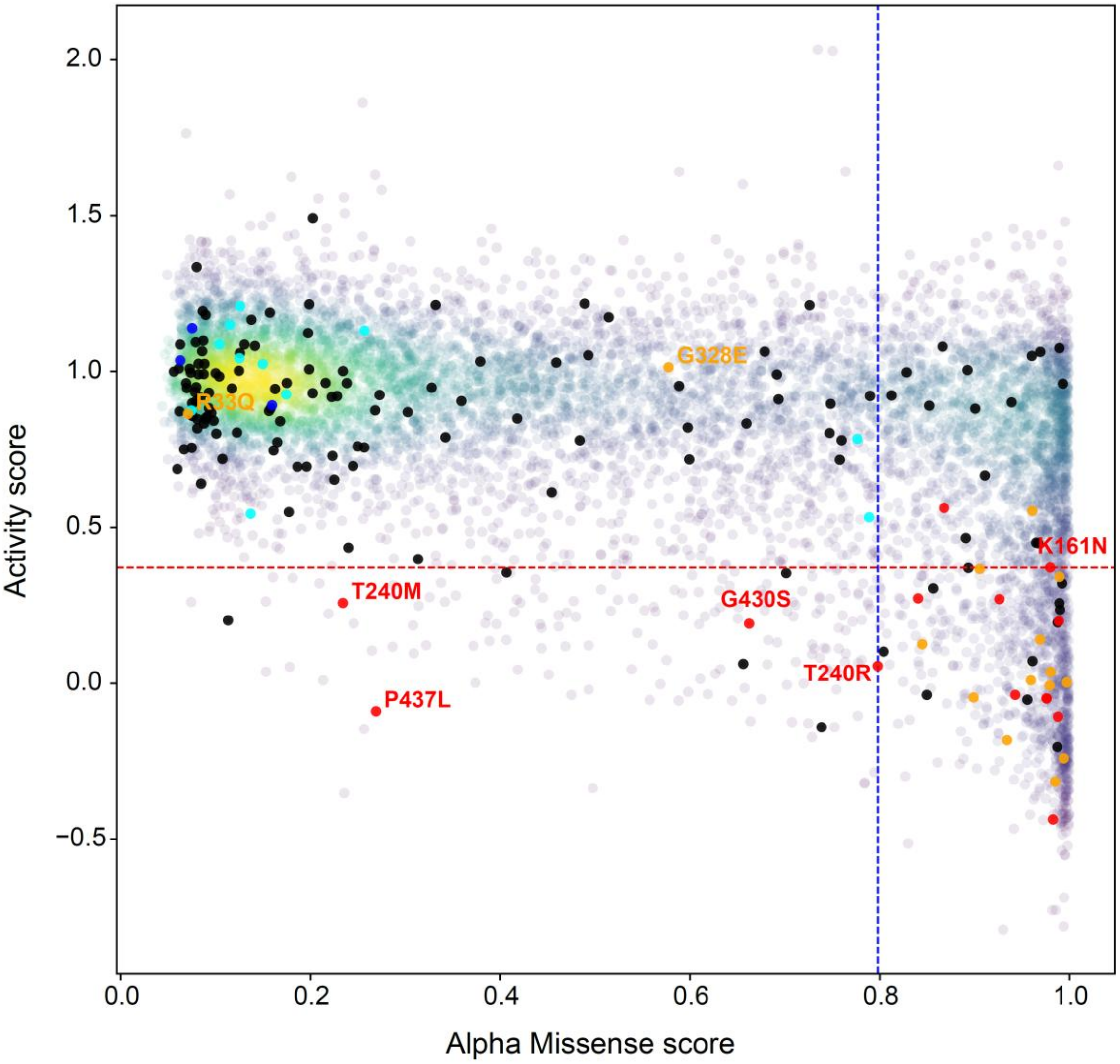


**Supplementary Figure 15.** *Activity and Alpha Missense predictions of pathogenic variants.* Scatterplot correlating Parkin activity scores with Alpha Missense scores for all missense Parkin variants. The known pathogenic (red) and likely pathogenic (orange) variants are labeled. Benign (blue), likely benign (cyan) variants, and variants of unknown significance (VUS, black) are highlighted in the plot. Blue and red dotted lines signify the thresholds as determined by Youden's J statistic that best separate pathogenic/likely pathogenic from benign/likely benign based on the Alpha Missense scores and activity scores respectively.

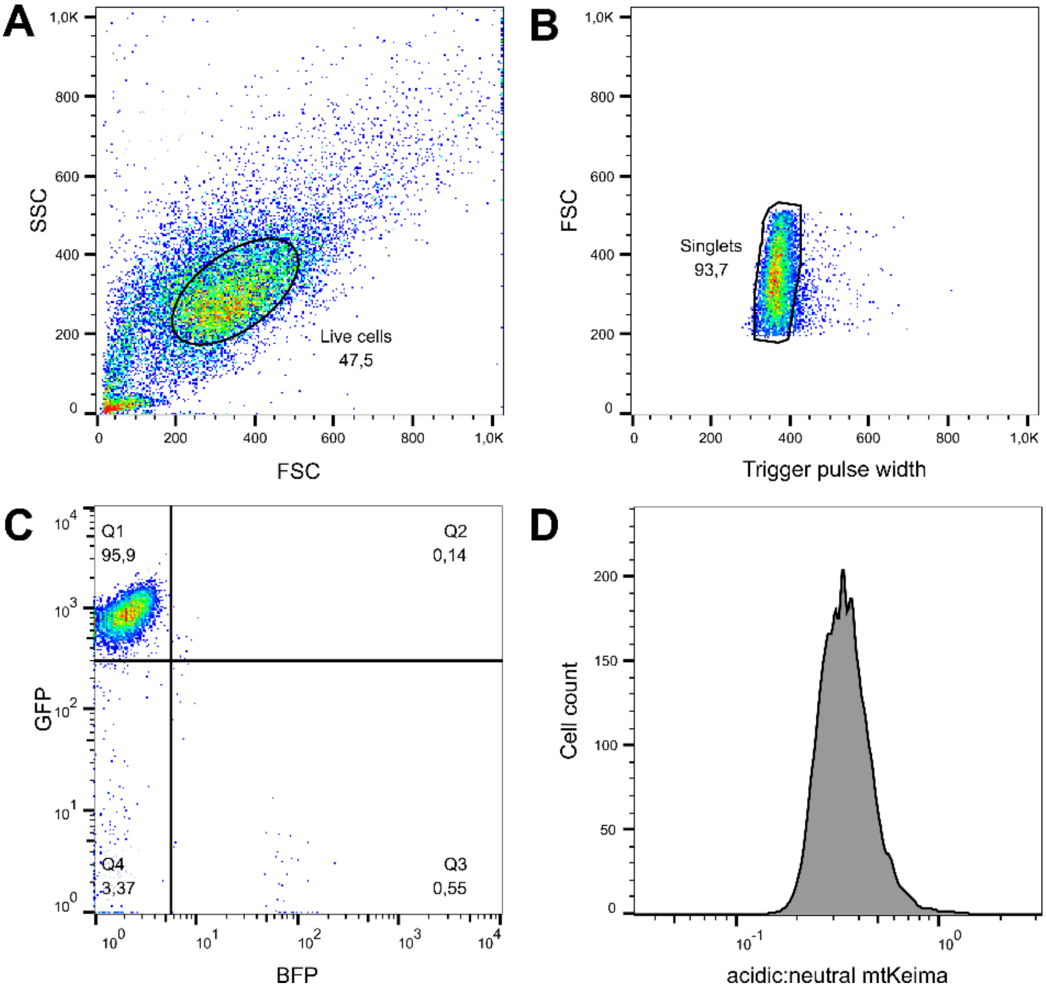


**Supplementary Figure 16.** *Gating strategy used for flow cytometry.* (A) Gating for live cells using forward scatter (FSC) and side scatter (SSC). (B) Singlets were selected based on trigger pulse width and FSC. (C) Successfully transfected cells were gated for GFP+/BFP-, corresponding to quadrant 1. (D) The histogram displays the ratio of acidic:neutral mtKeima to evaluate mitophagic activity. A high ratio indicates high mitophagic activity.

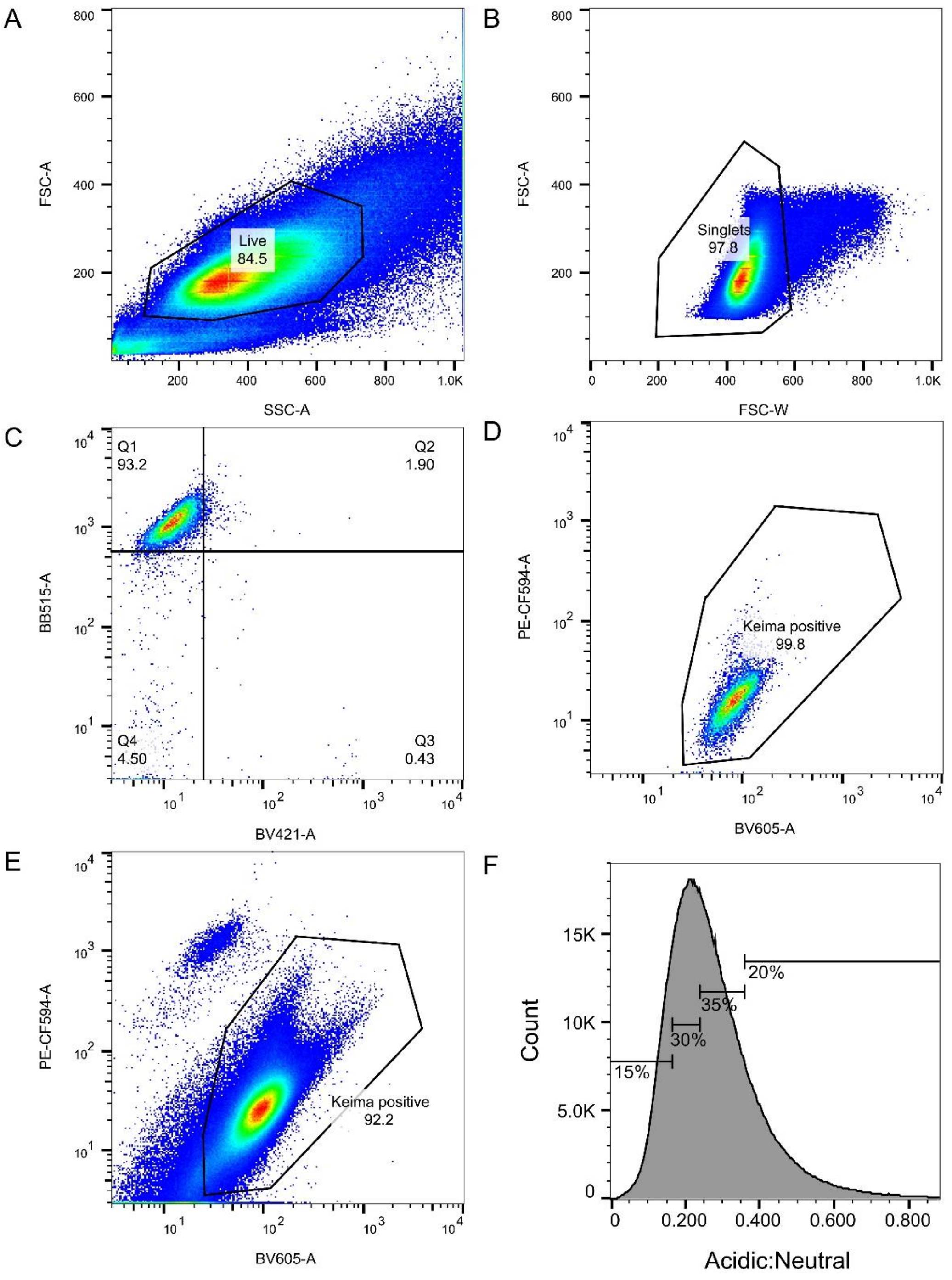


**Supplementary Figure 17.** *Full gating strategy for FACS of the Parkin variant library.* Parkin variant library transfected cells were gated for live (A) based on forward and side scatter, and (B) singlets, based on forward scatter area and forward scatter width. To gate for mtKeima positive cells the following strategy was used: (C) Flow cytometry data for cells transfected with the C431A Parkin variant was analysed (n=10,000) and the upper left quadrant (Q1) was found, corresponding to GFP positive and BFP negative cells. (D) Within that quadrant, the gate "Keima positive" was made, and this was then used for the library transfected singlets for the Parkin activity screen (E). Finally, a profile of the ratiometric of acidic:neutral mtKeima was plotted for the Keima positive cells and the total population was split into four bins covering 15%, 30%, 35% and 20% of the total cell population (F).

### Supplementary Table 1

*Variants selected for low throughput validation*

| Variant | Domain | Activity score | Abundance score[1] | Notes | Ref. |
|---|---|---|---|---|---|
| **WT** | - | 1.00 | 1.00 | Wild-type Parkin. Used for control and normalization. | - |
| **R42P** | UBL | 0.27 | -0.02 | Pathogenic. Potential hypomorph. Selected as a hypomorph control. | 2–4 |
| **A46T** | UBL | 0.53 | 0.65 | Within pUBL:RING0 binding interface. Selected for middle range activity. Benign. | 2,3,5 |
| **G47D** | UBL | 0.45 | 0.59 | Within pUBL:RING0 binding interface. Selected for middle range activity. | 5 |
| **Q57*** | UBL | -0.13 | 0.00 | Selected nonsense variant. | - |
| **S65A** | UBL | 0.26 | 0.99 | Site of PINK1 phosphorylation. Selected for low activity. | 4–6 |
| **V105G** | ACT | 0.59 | 0.98 | Important for ACT:RING0 binding and displacing RING2. Selected for middle range activity. | 5 |
| **K161N** | RING0 | 0.37 | 0.94 | Pathogenic. Key residue in pUBL:RING0 phospho-acceptor pocket. Selected for low activity. | 2,3,5 |
| **K211N** | RING0 | 0.34 | 1.01 | Pathogenic. Key residue in pUBL:RING0 phospho-acceptor pocket. Selected for low activity. | 2,3,5 |
| **V224A** | RING0 | 1.15 | 1.03 | Selected for hyperactivity. | 3,7,8 |
| **S246*** | RING1 | -0.25 | 0.04 | Selected nonsense variant. | - |
| **C253Y** | RING1 | 0.20 | -0.03 | Pathogenic and highly unstable. Selected for middle range activity. | 2,3 |
| **H302*** | Linker | -0.23 | -0.02 | Selected nonsense variant. | - |
| **R334C** | IBR | 1.02 | 0.48 | Selected for hyperactivity. | 3,6 |
| **T415N** | RING2 | 0.13 | 1.02 | Likely pathogenic. Residue implicated with catalytic activity. Selected for low activity. | 2,3,9 |
| **N428R** | RING2 | 0.51 | 1.13 | Close to the active site. Selected for middle range activity. | - |
| **G430D** | RING2 | -0.18 | 1.00 | In catalytic site. Pathogenic. Selected for low activity. | 2,3,6 |
| **C431A** | RING2 | 0.00 | 1.20 | Catalytic cystine residue. Used as negative control. | 10,11 |
| **E452D** | RING2 | 1.18 | 1.39 | Hyperstable. Selected for hyperactivity. | - |
| **V456A** | RING2 | 0.97 | 1.37 | Hyperstable. Selected for WT-like activity | - |

### Supplementary Table 2

*Notable positions in Parkin*

| Site | Domain | Med. activity score | Proposed mechanism | Ref. |
|---|---|---|---|---|
| **R6** | UBL | 0.72 | pUBL:RING0 interface (activated state) | [5,12,13] |
| **I44** | UBL | 0.46 | pUBL:RING0 interface (activated state) | [5,6,12–15] |
| **F45** | UBL | 0.33 | pUBL:RING0 interface (activated state) | [5,14] |
| **A46** | UBL | 0.45 | pUBL:RING0 interface (activated state) | [3,5,12] |
| **G47** | UBL | 0.46 | pUBL:RING0 interface (activated state) | [5] |
| **L61** | UBL | 0.38 | pUBL:RING0 interface (activated state) | [12] |
| **S65** | UBL | 0.16 | Activating phosphorylation site (PINK1) | [4–6,12,15–17] |
| **H68** | UBL | 0.72 | pUBL:RING0 interface (activated state) | [5,12,13] |
| **V70** | UBL | 0.44 | pUBL:RING0 interface (activated state) | [5,12,13] |
| **L102** | ACT | 0.62 | ACT (replaces M458 in RING2) | [1,5] |
| **V105** | ACT | 0.83 | ACT (replaces W462 in RING2) | [1,5] |
| **L107** | ACT | 0.50 | ACT (replaces F463 in RING2) | [1,5] |
| **Y143** | UBL-RING0 | 1.19 | Inhibitory phosphorylation site (c-Abl) | [8,18,19] |
| **C150** | RING0 | -0.20 | Zinc-coordination | [10] |
| **C154** | RING0 | 0.45 | Zinc-coordination | [10] |
| **K161** | RING0 | 0.26 | Phospho-acceptor pocket (pUBL:RING0 interface) | [5,6,13,14,17] |
| **R163** | RING0 | 0.15 | Phospho-acceptor pocket (pUBL:RING0 interface) | [5,6,13,14,17] |
| **V164** | RING0 | 0.48 | pUBL:RING0 interface (activated state) | [5] |
| **T173** | RING0 | 0.75 | pUBL:RING0 interface (activated state) | [5] |
| **T175** | RING0 | 0.39 | Activating phosphorylation site, pUBL:RING0 interface | [5,20,21] |
| **C201** | RING0 | 0.69 | Zinc-coordination | [10] |
| **K211** | RING0 | 0.15 | Phospho-acceptor pocket (pUBL:RING0 interface) | [5,6,13,14,17] |
| **C212** | RING0 | -0.14 | Zinc-coordination | [6,10] |
| **H215** | RING0 | 0.49 | Zinc-coordination | [10] |
| **T217** | RING0 | 0.22 | Activating phosphorylation site | [20,21] |
| **C238** | RING1 | -0.19 | Zinc-coordination | [10,21] |
| **T240** | RING1 | 0.06 | E2 binding site | [3,4,6,11,22,23] |
| **C241** | RING1 | -0.18 | Zinc-coordination | [10,21] |
| **D243** | RING1 | 0.69 | E2 binding site | [6,11] |
| **C253** | RING1 | 0.46 | Zinc-coordination | [10,21] |
| **H257** | RING1 | 0.62 | Zinc-coordination | [10,21] |
| **C260** | RING1 | -0.18 | Zinc-coordination | [10,21] |
| **C263** | RING1 | -0.20 | Zinc-coordination | [10,21] |
| **Y267** | RING1 | 0.46 | E2 binding site | [6] |
| **R275** | RING1 | 0.10 | pUb binding site | [3,12,13] |
| **G284** | RING1 | 0.55 | pUb binding site | [3,12,13,24] |
| **C289** | RING1 | -0.06 | Zinc-coordination | [6,10,21] |
| **V290** | RING1 | 0.68 | E2 binding site | [6] |
| **C293** | RING1 | 0.74 | Zinc-coordination | [10,21] |

| | | | | |
|---|---|---|---|---|
| **H302** | RING1 | 0.42 | pUb binding site | 5,12,13,17,24,25 |
| **G319** | IBR | 0.37 | pUb binding site | 24 |
| **A320** | IBR | 0.69 | pUb binding site | 22,24 |
| **E321** | IBR | 0.23 | pUb binding site | 3,14,15 |
| **V324** | IBR | 0.81 | pUb binding site | 24 |
| **C332** | IBR | -0.13 | Zinc-coordination | 10,21 |
| **C337** | IBR | -0.14 | Zinc-coordination | 10,21 |
| **G340** | IBR | 0.26 | pUb binding site | 24 |
| **C352** | IBR | -0.14 | Zinc-coordination | 10,21 |
| **C360** | IBR | 0.19 | Zinc-coordination | 10,21 |
| **C365** | IBR | 0.57 | Zinc-coordination | 10,21 |
| **C368** | IBR | 0.52 | Zinc-coordination | 10,21 |
| **S407** | IBR-RING2 | 0.14 | Linker interacts with donor Ub from E2 | 9,17 |
| **T410** | IBR-RING2 | 0.59 | Linker interacts with donor Ub from E2 | 9,17 |
| **I411** | IBR-RING2 | 0.10 | Linker interacts with donor Ub from E2 | 9,17 |
| **T414** | IBR-RING2 | 0.59 | Linker interacts with donor Ub from E2 | 17 |
| **T415** | IBR-RING2 | 0.12 | Linker interacts with donor Ub from E2 | 3,9,17 |
| **K416** | IBR-RING2 | 0.33 | Linker interacts with donor Ub from E2 | 17 |
| **C418** | RING2 | -0.31 | Zinc-coordination | 10,21,26 |
| **C421** | RING2 | -0.29 | Zinc-coordination | 10,21 |
| **E426** | RING2 | 0.11 | Transfer of donor Ub | 9 |
| **G430** | RING2 | 0.10 | Transfer of donor Ub | 3,4,6 |
| **C431** | RING2 | 0.00 | Catalytic site, thioester bond to Ub | 4,6,10,11,13,23 |
| **H433** | RING2 | 0.46 | Catalytic site | 6,8,10,11,23,27 |
| **C436** | RING2 | -0.33 | Zinc-coordination | 10,21 |
| **P437** | RING2 | 0.39 | Transfer of donor Ub | 9 |
| **C441** | RING2 | -0.25 | Zinc-coordination | 6,10,21 |
| **C446** | RING2 | 0.07 | Zinc-coordination | 10,21 |
| **C449** | RING2 | 0.33 | Zinc-coordination | 10,21 |
| **C451** | RING2 | 1.34 | Inhibitory succination site | 28 |
| **M458** | RING2 | 1.24 | RING0:RING2 interface (autoinhibited state) | 3,5 |
| **H461** | RING2 | 0.74 | Zinc-coordination | 10,21 |
| **W462** | RING2 | 1.07 | RING0:RING2 interface (autoinhibited state) | 5,27,29 |
| **D464** | RING2 | 1.05 | RING0:RING2 interface (autoinhibited state) | 5 |

### Supplementary Table 3

*Positions without known mechanism*

| Site | Domain | Med. activity score | Notes |
|---|---|---|---|
| **V3** | UBL | 0.45 | Likely in the pUBL:RING0 interface (R6 is[5]) |
| **V5** | UBL | 0.52 | Likely in the pUBL:RING0 interface (R6 is[5]) |
| **V15** | UBL | 0.54 | Structurally important[1] |
| **V17** | UBL | 0.65 | Structurally important[1] |
| **I23** | UBL | 0.45 | Structurally important[1] |
| **L26** | UBL | 0.41 | Structurally important[1] |
| **K27** | UBL | 0.47 | Structurally important[1] |
| **V30** | UBL | 0.51 | Structurally important[1] |
| **A31** | UBL | 0.55 | Structurally important[1] |
| **L41** | UBL | 0.58 | Likely in the pUBL:RING0 interface (I44 hydrophobic patch[5]) |
| **V43** | UBL | 0.55 | Likely in the pUBL:RING0 interface (I44 hydrophobic patch[5]) |
| **L50** | UBL | 0.49 | Likely in the pUBL:RING0 interface (I44 hydrophobic patch[5]) |
| **V56** | UBL | 0.54 | Likely in the pUBL:RING0 interface (I44 hydrophobic patch[5]) |
| **C59** | UBL | 0.54 | Likely in the pUBL:RING0 interface (area around S65[5]) |
| **V67** | UBL | 0.43 | Likely in the pUBL:RING0 interface (area around S65[5]) |
| **I69** | UBL | 0.50 | Likely in the pUBL:RING0 interface (area around S65[5]) |
| **S141** | UBL:RING0 | 1.05 | Hyperactivity patch. Likely in the RING0:RING interface (146 is[5]) |
| **V148** | RING0 | 0.80 | Structurally important[1] |
| **R156** | RING0 | 1.08 | Hyperactivity patch. Likely in the RING0:RING2 interface (162 is[5]) |
| **S223** | RING0 | 0.62 | Unknown |
| **S233** | RING0-RING1 | 1.08 | Undefined hyperactivity |
| **P294** | RING1-IBR | 1.08 | Undefined hyperactivity |
| **F304** | RING1-IBR | 0.77 | Likely part of the pUb binding site. (R305 is[5]) |
| **R314** | IBR | 0.52 | Structurally important[1] |
| **G329** | IBR | 0.00 | Structurally important[1] |
| **P419** | RING2 | 0.48 | Close to active site |
| **V423** | RING2 | 0.16 | Close to active site |
| **V425** | RING2 | 0.17 | Close to active site |
| **K427** | RING2 | 0.15 | Close to active site |
| **G429** | RING2 | 0.58 | Close to active site |
| **M434** | RING2 | 0.01 | Close to active site |
| **W445** | RING2 | -0.19 | Structurally important[1] |
| **W447** | RING2 | -0.02 | Unknown |
| **E452** | RING2 | 1.15 | Unknown hyperactivity |

### Supplementary Table 4

*Notable positions where substitutions cause minor effects*

| Site | Domain | Med. activity score | Proposed mechanism | Ref. |
|---|---|---|---|---|
| **C166** | RING0 | 0.93 | Zinc-coordination | [10] |
| **C169** | RING0 | 0.92 | Zinc-coordination | [10] |
| **C196** | RING0 | 0.93 | Zinc-coordination | [10] |
| **H373** | IBR | 0.85 | Zinc-coordination | [10,21] |
| **C377** | IBR | 0.90 | Zinc-coordination | [10,21] |
| **C457** | RING2 | 0.82 | Zinc-coordination | [10,14,21] |
| **S108** | ACT | 0.82 | ULK1 phosphorylation site | [30] |
| **K151** | RING0 | 0.80 | pUb binding site | [5,12,17,24] |
| **R305** | RING1-IBR | 0.96 | pUb binding site | [5,12,13,17] |
| **F146** | RING0 | 0.81 | RING0:RING2 interface (autoinhibited state) | [3,5,8,14] |
| **L162** | RING0 | 0.91 | RING0:RING2 interface (autoinhibited state) | [5] |
| **P180** | RING0 | 0.88 | RING0:RING2 interface (autoinhibited state) | [5] |
| **W183** | RING0 | 0.72 | RING0:RING2 interface (autoinhibited state) | [5] |
| **V186** | RING0 | 0.92 | RING0:RING2 interface (autoinhibited state) | [5] |
| **F208** | RING0 | 0.89 | RING0:RING2 interface (autoinhibited state) | [5,8] |
| **F210** | RING0 | 0.94 | RING0:RING2 interface (autoinhibited state) | [5] |
| **F463** | RING2 | 0.91 | RING0:RING2 interface (autoinhibited state) | [5,14] |
| **V465** | RING2 | 0.99 | RING0:RING2 interface (autoinhibited state) | [5] |
| **E444** | RING2 | 0.84 | Catalytic triad | [6,11,23,27] |

### Supplementary Table 5

*Primers used in this study*

| Primer | Sequence |
|---|---|
| LC1020 | CCAGGACATATGAGGACTAG |
| LC1031 | GGGTTAGCAAGTGGCAGCCTTCTCCTTAATCAGCTCTTCG |
| JS_R | CAAGCAGAAGACGGCATACGAGAT (NNNNNNNN) GGGTTAGCAAGTGGCAGCCT |
| PCR2_Fw1 | AATGATACGGCGACCACCGAGATCTACAC (TAGCAGTC) CCAGGACATATGAGGACTAG |
| LC1040 | AAGAACCGCTAGAAGCGTCGCTGTACAAATAGTT |
| LC1041 | CGAGAAAGCTAGCGCAAACGACTACTCGCA |
| LC1042 | ctgattaaggagaAGGCTGCCACTTGCTAACCC |
| ASPA_PARK2_index2_re | ACGCAATTGCAGAACTAGTCCTCATATGTCCTGG |
| VV230 | GGTTGTGGCCATATTATCATCGTGTTT |
| VV231 | GGATCCTGATCATAATCAGCCATACCA |
| VV232 | ATGTGGTATGGCTGATTATGATCAGGATCC |
| VV233 | GAAAAACACGATGATAATATGGCCACAACC |
